# Emergence of brain-like mirror-symmetric viewpoint tuning in convolutional neural networks

**DOI:** 10.1101/2023.01.05.522909

**Authors:** Amirhossein Farzmahdi, Wilbert Zarco, Winrich Freiwald, Nikolaus Kriegeskorte, Tal Golan

## Abstract

Primates can recognize objects despite 3D geometric variations such as in-depth rotations. The computational mechanisms that give rise to such invariances are yet to be fully understood. A curious case of partial invariance occurs in the macaque face-patch AL and in fully connected layers of deep convolutional networks in which neurons respond similarly to mirror-symmetric views (e.g., left and right profiles). Why does this tuning develop? Here, we propose a simple learning-driven explanation for mirror-symmetric viewpoint tuning. We show that mirror-symmetric viewpoint tuning for faces emerges in the fully connected layers of convolutional deep neural networks trained on object recognition tasks, even when the training dataset does not include faces. First, using 3D objects rendered from multiple views as test stimuli, we demonstrate that mirror-symmetric viewpoint tuning in convolutional neural network models is not unique to faces: it emerges for multiple object categories with bilateral symmetry. Second, we show why this invariance emerges in the models. Learning to discriminate among bilaterally symmetric object categories induces reflection-equivariant intermediate representations. AL-like mirror-symmetric tuning is achieved when such equivariant responses are spatially pooled by downstream units with sufficiently large receptive fields. These results explain how mirror-symmetric viewpoint tuning can emerge in neural networks, providing a theory of how they might emerge in the primate brain. Our theory predicts that mirror-symmetric viewpoint tuning can emerge as a consequence of exposure to bilaterally symmetric objects beyond the category of faces, and that it can generalize beyond previously experienced object categories.

## Introduction

Primates can recognize objects robustly despite considerable image variation. Although we experience object recognition as immediate and effortless, the process involves a large portion of cortex and considerable metabolic cost, and determining the neural mechanisms and computational principles that enable this ability remains a major neuroscientific challenge. One particular object category, faces, offers an especially useful window into how the visual cortex transforms retinal signals to object representations. The macaque brain contains a network of interconnected areas devoted to the processing of faces. This network, the face-patch system, forms a subsystem of the inferotemporal (IT) cortex [1– Neurons across the network show response selectivity for faces, but are organized in face patches–spatially and functionally distinct modules [3, 5]. These patches exhibit an information processing hierarchy from posterior to anterior areas. In the most posterior face-patch, PL (posterior lateral), neurons respond to face components [6]. In ML/MF (middle lateral/middle fundus), neurons respond to whole faces in a view-specific manner. In AL (anterior lateral), responses are still view-specific, but mostly reflectioninvariant. Finally in AM (anterior medial), neurons respond with sensitivity to the identity of the face, but in a view-invariant fashion [3]. The average neuronal response latencies increase across this particular sequence of stages [3]. Thus, it appears as if visual information is transformed across this hierarchy of representational stages in a way that facilitates the recognition of faces despite view variations.

What are the computational principles that give rise to the representational hierarchy evident in the face-patch system? Seeking potential answers to this and similar questions, neuroscientists have been increasingly turning to convolutional neural networks (CNNs) as initial computational models of primate ventral stream vision. While CNNs lack many seemingly essential biological details, they offer a simple hierarchical model of the cascade of linear-non-linear transformations carried out by the feedforward sweep of computations in the ventral visual stream. Feedforward CNNs remain among the best models for predicting mid- and high-level cortical representations of novel natural images within the first 100-200 ms after stimulus onset [7, 8]. Diverse CNN models, trained on tasks such as face identification [9–1 object recognition [12], inverse graphics [13], sparse coding [14], and unsupervised generative modeling [15] have all been shown to replicate at least some aspects of face-patch system representations. Face-selective artificial neurons occur even in untrained CNNs [16], and functional specialization between object and face representation emerges in CNNs trained on the dual task of recognizing objects and identifying faces [17].

To better characterize and understand the computational mechanisms employed by the primate face-patch system and test whether the assumptions implemented by current CNN models are sufficient for explaining its function, we should carefully inspect the particular representational motifs the face-patch system exhibits. One of the more salient and intriguing of these representational motifs is the *mirror-symmetric viewpoint tuning* in the AL face-patch [3]. Neurons in this region typically respond with different firing rates to varying views of a face (e.g., a lateral profile vs. a frontal view), but they respond with similar firing rates to views that are horizontal reflections of each other (e.g., left and right lateral profiles) [3].

To date, two distinct computational models have been put forward as potential explanations for AL’s mirror-symmetric viewpoint tuning. Leibo and colleagues [18] considered unsupervised learning in an HMAX-like [19] four-layer neural network exposed to a sequence of face images rotating in depth about a vertical axis. When the learning of the mapping from the complex-cell-like representation of the second layer to the penultimate layer was governed by Hebbian-like synaptic updates (Oja’s rule, [20]), approximating a principal components analysis of the input images, the penultimate layer developed mirror-symmetric viewpoint tuning. In another modeling study, Yildirim and colleagues [13] trained a CNN to invert the rendering process of 3D faces, yielding a hierarchy of intermediate and high-level face representations. Mirror-symmetric viewpoint tuning emerged in an intermediate representation between two densely-connected transformations mapping 2.5D surface representations to high-level shape and texture face-space representations. Each of these two models [13, 18] provides a plausible explanation of AL’s mirror-symmetric viewpoint tuning, but each requires particular assumptions about the architecture and learning conditions, raising the question whether a more general computational principle can provide a unifying account of the emergence of mirror-symmetric viewpoint tuning.

Here, we propose a parsimonious, bottom-up explanation for the emergence of mirror-symmetric viewpoint tuning for faces (Fig. 1). We find that learning to discriminate among bilaterally symmetric object categories promotes the learning of representations that are *reflection-equivariant* (i.e., they code a mirror image by a mirrored representation). Spatial pooling of the features, as occurs in the transition between the convolutional and fully connected layers in AlexNet or VGG16-like CNNs, then yields *reflection-invariant* representations (i.e., these representations code a mirror image as they would code the original image). These reflection-invariant representations are not fully view-invariant: They are still tuned to particular views of faces (e.g., respond more to a half-profile than to a frontal view, or vice versa), but they do not discriminate between mirrored views. In other words, these representations exhibit mirror-symmetric viewpoint tuning (in the twin sense of the neuron responding equally to left-right-reflected images and the tuning function, hence, being mirror-symmetric). We propose that the same computational principles may explain the emergence of mirror-symmetric viewpoint tuning in the primate face-patch system.

**Figure 1.**
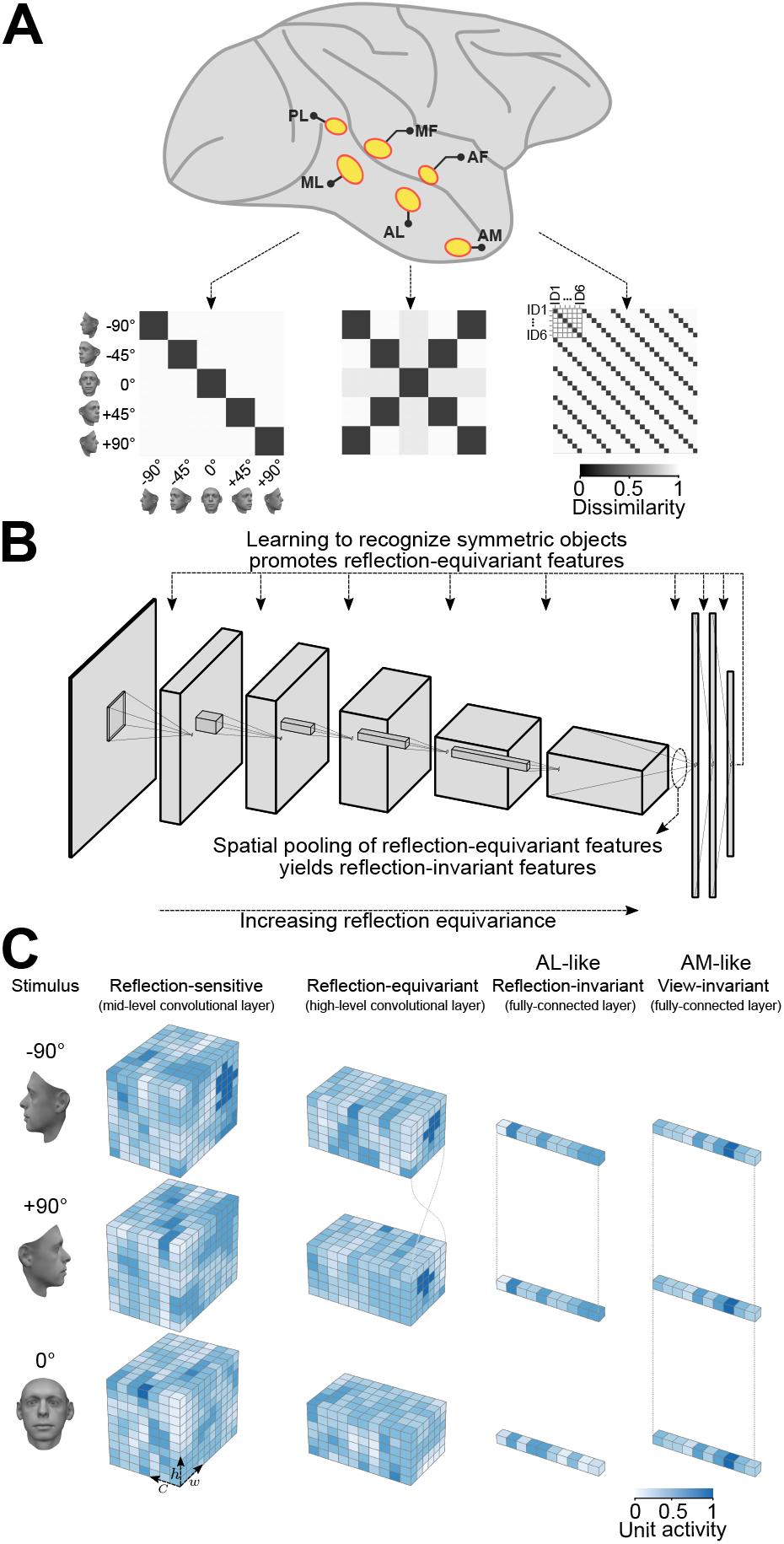
An overview of our claim: convolutional deep neural networks trained on discriminating among bilaterally symmetric object categories provide a parsimonious explanation for the mirror-symmetric viewpoint tuning of the macaque AL face-patch. (**A**) The macaque face-patch system. Face-selective cortical areas are highlighted in yellow. The areas ML, AL, and AM exhibit substantially different tuning proprieties when presented with faces of different head orientations [3]. We illustrate this empirical finding here by schematic population-level representational dissimilarity matrices (RDMs). From posterior to anterior face areas, invariance to viewpoints gradually increases: from view-tuned in ML, through mirror-symmetric in AL, to view-invariant identity selectivity in AM. (**B**) Training convolutional deep neural networks on recognizing specific symmetric object categories (e.g., faces, cars, the digit 8) gives rise to AL-like mirror-symmetric tuning. It is due to a cascade of two effects: First, learning to discriminate among symmetric object categories promotes tuning for reflection-equivariant representations throughout the entire processing layers. This reflection equivariance increases with depth. Then, long-range spatial pooling (as in the transformation of the last convolution layer to the first fully connected layer in CNNs) transforms the equivariant representations into reflection invariant representations. (**C**) Schematic representations of three viewpoints of a face (left profile, frontal view, right profile) are shown in three distinct stages of processing. Each tensor depicts the width (w), height (h), and depth (c) of an activation pattern. Colors indicate channel activity. From left to right: In a mid-level convolutional layer, representations are view-specific. A deeper convolutional layer produces reflection-equivariant representations that are view-specific. Feature vectors of a fully connected layer become invariant to reflection by pooling reflection-equivariant representations from the last convolutional layer.

Our results further suggest that emergent reflection-invariant representations may also exist for non-face objects: the same training conditions give rise to CNN units that show mirror-symmetric tuning profiles for non-face objects that have a bilaterally symmetric structure. Extrapolating from CNNs back to primate brains, we predict AL-like mirror-symmetric viewpoint tuning in non-face-specific visual regions that are parallel to AL in terms of the ventral stream representational hierarchy. Such tuning could be revealed by probing these regions with non-face objects that are bilaterally symmetric.

## Results

### Deep layers in CNNs exhibit mirror-symmetric view-point tuning to multiple object categories

We investigated whether reflection-invariant yet view-specific tuning emerges naturally in deep convolutional neural networks. To achieve this, we generated a diverse set of 3D objects rendered in multiple views. Different 3D object categories have different numbers of symmetry planes. For example, a face has a single approximate symmetry plane: its left and right halves are reflected versions of each other. A car viewed from the outside typically also has a symmetry plane dividing the left and right halves. In addition, some car models have an approximate front-back symmetry, with the front and back wheels and the hood and trunk approximately matching. However, a flower typically does not have a clear symmetry plane (Fig. 2A), although it may have radial symmetry. We evaluated the hidden-layer activations of an ImageNet-trained AlexNet CNN model [21] presented with nine views of each object exemplar. We constructed a 9 ×9 representational dissimilarity matrix (RDM, [22]) for each exemplar object and each CNN layer, summarizing the view tuning of the layer’s artificial neurons (“units”) by means of between-view representational distances. The resulting RDMs revealed a progression throughout the CNN layers for objects with one or more symmetry planes: These objects induce mirror-symmetric RDMs in the deeper CNN layers (Fig. 2B), reminiscent of the symmetric RDMs measured for face-related responses in the macaque AL face-patch [3]. We defined a “mirror-symmetric viewpoint tuning index” to quantify the degree to which representations are view-selective yet reflection-invariant (Fig. 2C). Consider a dissimilarity matrix *D ∈* ℝ^*n×n*^ where *D*_*j,k*_ denotes the distance between view *j* and view *k, n* denotes the number of views. The RDM is symmetric about the main diagonal by definition: *D*_*j,k*_ = *D*_*k,j*_, independent of the tuning of the units. The views are ordered from left to right, such that *j* and *n* + 1 − *k* refer to horizontally reflected views. The mirror-symmetric view-point tuning index is defined as the Pearson linear correlation coefficient between *D* and its horizontally flipped counterpart, 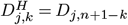 (Eq. 1). Note that this is equivalent to the correlation between vertically flipped RDMs, because of the symmetry of the RDMs about the diagonal: 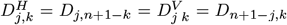. This mirror-symmetric viewpoint tuning index is positive and large to the extent that the units are view-selective but reflection-invariant (like the neurons in macaque AL face-patch). The index is near zero for units with view-invariant tuning (such as the AM face-patch), where the dissimilarities are all small and any variations are caused by noise.

**Figure 2.**
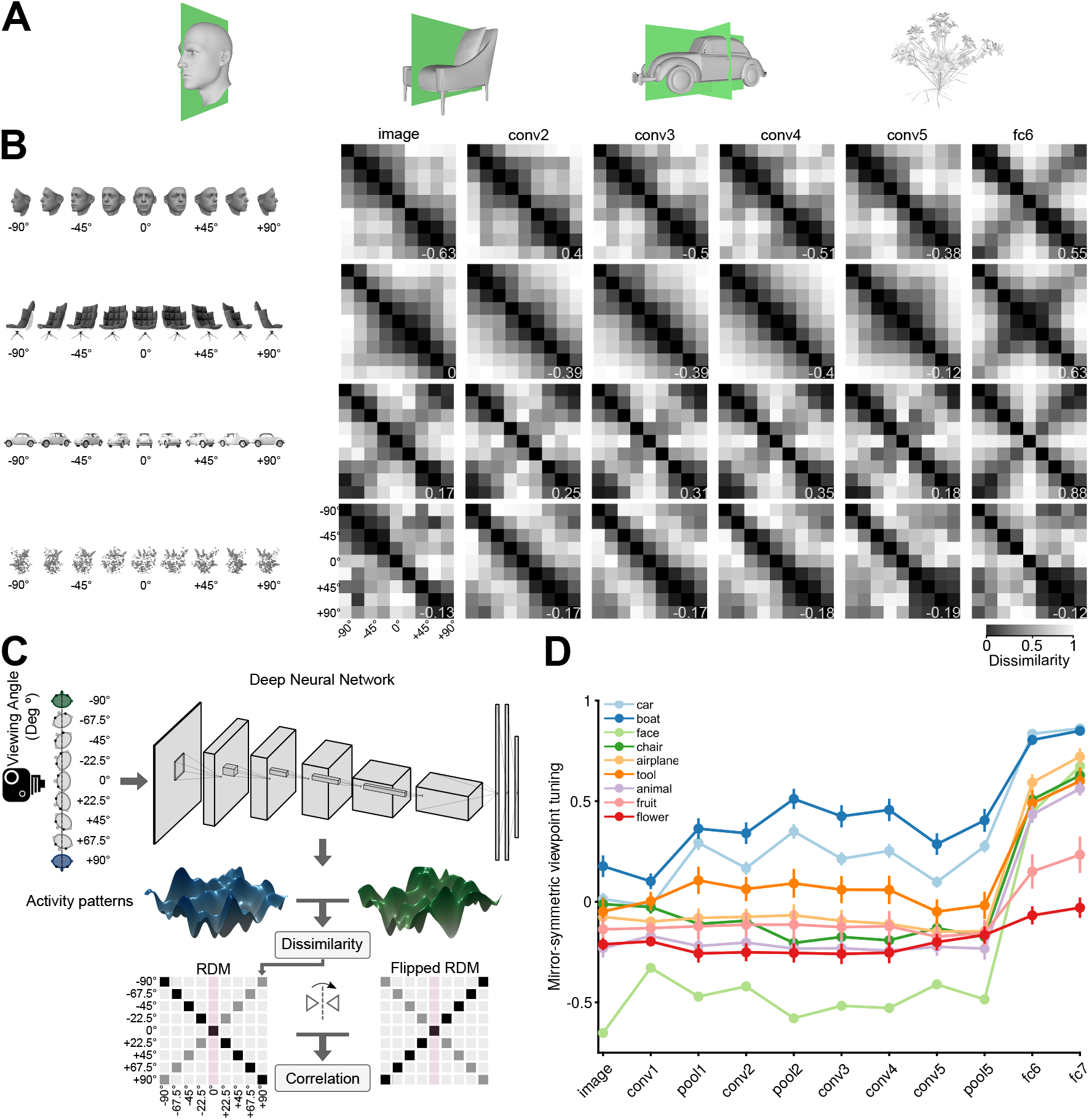
Mirror-symmetric viewpoint tuning of higher-level deep neural network representations emerges for multiple object categories. (**A**) 3D Objects have different numbers of symmetry axes. (**B**) Different viewpoint tuning across the layers of AlexNet for four example objects. A face (top row), a non-face object with bilateral symmetry (a chair, second row), an object with quadrilateral symmetry (a car, third row), and an object with no obvious reflective symmetry planes (a flower, bottom row). For each object, the responses to nine views (−90°to +90°in the steps of 22.5°) were measured in six key AlexNet layers, shallow (input, *left*) to deep (fc6, *right*). For each layer, a Representational Dissimilarity Matrix (RDM) depicts how the population activity vector varies across different object views. Each element of the RDM represents the dissimilarity (1 *−* Pearson correlation coefficient) between a pair of activity vectors evoked in response to two particular views. The symmetry of the RDMs about the major diagonal is inherent to their construction. However, the symmetry about the minor diagonal (for the face and chair, in fc6, and for the car, already in conv2) indicates mirror-symmetric viewpoint tuning. (**C**) The schematic shows how the mirror-symmetric viewpoint tuning index was quantified. We first fed the network with images of each object from nine viewpoints and recorded the activity patterns of its layers. Then, we computed the dissimilarity between activity patterns of different viewpoints to create an RDM. Next, we measured the correlation between the obtained RDM and its horizontally flipped counterpart, excluding the frontal view (which is unaffected by the reflection). (**D**) The Mirror-symmetric viewpoint tuning index across all AlexNet layers for nine object categories (car, boat, face, chair, airplane, animal, tool, fruit, and flower). Each solid circle denotes the average of the index over 25 exemplars within each object category. Error bars indicate the standard error of the mean. The mirror-symmetric viewpoint tuning index values of the four example objects in panel B are shown at the bottom right of each RDM in panel B.

Fig. 2D shows the average mirror-symmetric viewpoint tuning index for each object category across AlexNet layers. For object categories with a bilaterally symmetric structure (faces, chairs, airplanes, tools, and animals), the mirror-symmetric viewpoint tuning index throughout the convolutional layers was low (less than 0.1) or even negative. Once the signal arrived at the first fully connected layer (fc6), the score abruptly leaped to high values (greater than 0.6). By contrast, fruits and flowers, two categories without apparent symmetry planes, induced low index values both in the convolutional and the fully connected layers. Cars and boats, two categories with left-right as well as front-back symmetry structure, induced high mirror-symmetric viewpoint tuning index levels throughout the layers. For cars and boats, the additional symmetry plane causes reflected images of the profile views (where the view axis is parallel to the second symmetry plane) to be similar, even at the pixel level, which elevates the mirror-symmetric viewpoint tuning index already in shallow layers.

Convolutional neural networks of a different architecture (VGG16, [23]) and a different objective function (face identification, [24]) reproduce this pattern of results (Fig. S1). However, for an untrained AlexNet, the mirror-symmetric viewpoint tuning index remains relatively constant across the layers (Fig. S2A). Statistically contrasting mirror-symmetric viewpoint tuning between a trained and untrained AlexNet demonstrates that the leap in mirror-symmetric viewpoint tuning in fc6 is training-dependent (Fig. S2B).

Why does the transition to the fully connected layers induce mirror-symmetric viewpoint tuning for bilaterally symmetric objects? One potential explanation is that the learned weights that map the last convolutional representation (pool5) to the first fully connected layer (fc6) combine the pool5 activations in a specific pattern that induces mirror-symmetric viewpoint tuning. However, replacing fc6 with spatial global average pooling (collapsing each pool5 feature map into a scalar activation) yields a representation with very similar mirror-symmetric viewpoint tuning levels (Fig. S3). This result is suggestive of an alternative explanation: that training the network on ImageNet gives rise to a reflection-equivariant representation in pool5. We therefore investigated the reflection equivariance of the convolutional representations.

### Reflection equivariance versus reflection invariance of convolutional layers

Consider a representation *f* (·), defined as a function that maps input images to sets of feature maps, and a geometric image transformation *g*(·), applicable to either feature maps or raw images. *f* is equivariant under *g* if *f* (*g*(*x*)) = *g*(*f* (*x*)) for any input image *x* (see also [25]). While convolutional feature maps are approximately equivariant under translation (but see [26]), they are not in general equivariant under reflection or rotation. For example, an asymmetrical filter along reflection axes in the first convolutional layer would yield an activation map that is not equivariant under reflection. And yet, the demands of the task on which a CNN is trained may lead to the emergence of representations that are approximately equivariant under reflection or rotation (see [27, 28] for neural network architectures that are equivariant to reflection or rotation by construction).

If a representation *f* is equivariant under a transformation *g* that is a spatial permutation of its input (e.g., *g* is a horizontal or vertical reflection or a 90° rotation) then *f* (*x*) and *f* (*g*(*x*)) are spatially permuted versions of each other. If a spatially invariant function *h*(·) (i.e., a function that treats the pixels as a set, such as the average or the maximum) is then applied to the feature maps, the composed function *h ° f* is *invariant* to *g* since *h (f* (*g*(*x*))) = *h (g*(*f* (*x*))) = *h*(*f* (*x*)). Transforming a stack of feature maps into a channel vector by means of global average pooling is a simple case of such a spatially invariant function *h*. Therefore, if task-training induces approximately reflection-equivariant representations in the deepest convolutional layer of a CNN and approximately uniformly pooling in the following fully connected layer, the resulting pooled representation would be approximately reflection-invariant.

We examined the emergence of approximate equivariance and invariance in CNN layers (Fig. 3). We considered three geometric transformations: horizontal reflection, vertical reflection, and 90° rotation. Note that given their architecture alone, CNNs are not expected to show greater equivariance and invariance for horizontal reflection compared to vertical reflection or 90° rotation. However, greater invariance and equivariance for horizontal reflection may be expected on the basis of natural image statistics and the demands of invariant recognition. Many object categories in the natural world are bilaterally symmetric with respect to a plane parallel to the axis of gravity and are typically viewed (or photographed) in an upright orientation. Horizontal image reflection, thus, tends to yield equally natural images of similar semantic content, whereas vertical reflection and 90°rotation yield unnatural images.

**Figure 3.**
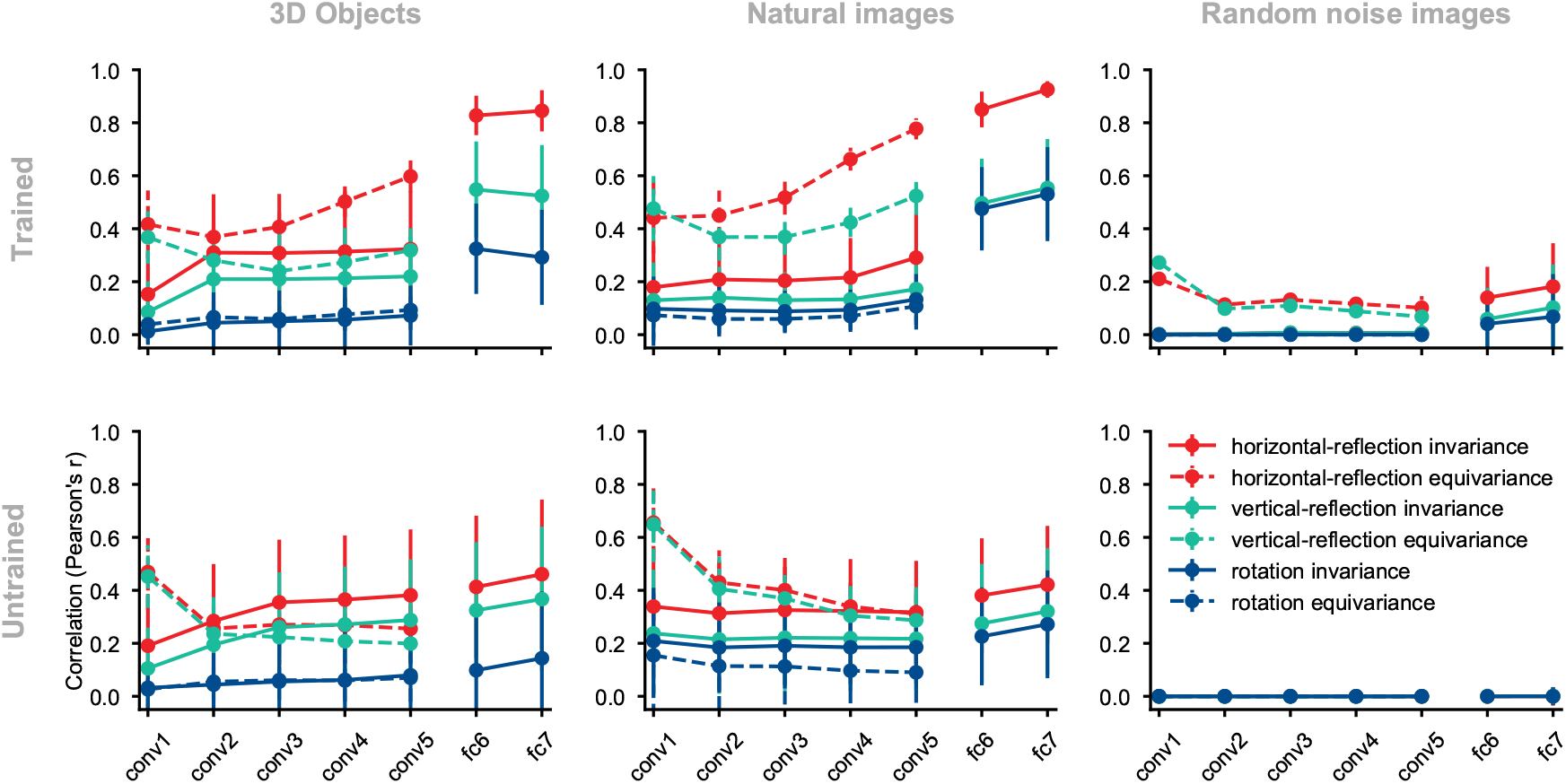
Equivariance and invariance in trained and untrained deep convolutional neural networks. Each solid circle represents an equivariance or invariance measure, averaged across images. Hues denote different transformations (horizontal flipping, vertical flipping, or 90° rotation). Error bars depict the standard deviation across images (each test condition consists of 2025 images). Invariance is a measure of similarity between the activity pattern an image elicits and the activity pattern its transformed (e.g., flipped) counterpart (solid lines) elicits. Equivariance is a measure of the similarity between the activity pattern of a transformed image elicits and the *transformed* version of the activity pattern the untransformed image elicits (dashed lines). (**A**) ImageNet-trained AlexNet tested on the rendered 3D objects. (**B**) Untrained AlexNet tested on rendered 3D objects. (**C**) ImageNet-trained AlexNet tested on the natural images (images randomly selected from the test set of ImageNet). (**D**) Untrained AlexNet tested on the natural images. (**E**) ImageNet-trained AlexNet tested on the random noise images. (**F**) Untrained AlexNet tested on the random noise images.

To measure equivariance and invariance, we presented the CNNs with pairs of original and transformed images. To measure the invariance of a fully-connected CNN layer, we calculated an across-unit Pearson correlation coefficient for each pair of activation vectors that were induced by a given image and its transformed version. We averaged the resulting correlation coefficients across all image pairs (Materials and Methods, Eq. 2). For convolutional layers, this measure was applied after flattening stacks of convolutional maps into vectors. In the case of horizontal reflection, this invariance measure would equal 1.0 if the activation vectors induced by each image and its mirrored version are identical (or perfectly correlated).

Equivariance could be quantified only in convolutional layers because units in fully connected layers do not form visuotopic maps that can undergo the same transformations as images. It was quantified similarly to invariance, except that we applied the transformation of interest (i.e., reflection or rotation) not only to the image but also to the convolutional map of activity elicited by the untransformed image (Eq 3). We correlated the representation of the transformed image with the transformed representation of the image. In the case of horizontal reflection, this equivariance measure would equal 1.0 if each activation map induced by an image and its reflected version are reflected versions of each other (or are perfectly correlated after horizontally flipping one of them).

We first evaluated equivariance and invariance with respect to the set of 3D object images described in the previous section. In an ImageNet-trained AlexNet, horizontal-reflection equivariance increased across convolutional layers (Fig. 3A). Equivariance under vertical reflection was less pronounced and equivariance under 90°rotation was even weaker (Fig. 3A). In this trained AlexNet, invariance jumped from a low level in convolutional layers to a high level in the fully connected layers and was highest for horizontal reflection, lower for vertical reflection, and lowest for 90°rotation.

In an untrained AlexNet, the reflection equivariance of the first convolutional layer was higher than in the trained network. However, this measure subsequently decreased in the deeper convolutional layers to a level lower than that observed for the corresponding layers in the trained network. The higher level of reflection-equivariance of the first layer of the untrained network can be explained by the lack of strongly oriented filters in the randomly initialized layer weights. While the training leads to oriented filters in the first layer, it also promotes downstream convolutional representations that have greater reflection-equivariance than those in a randomly-initialized, untrained network.

The gap between horizontal reflection and vertical reflection in terms of both equivariance and invariance was less pronounced in the untrained network (Fig. 3B), indicating a contribution of task training to the special status of horizontal reflection. In contrast, the gap between vertical reflection and 90° rotation in terms of both equivariance and invariance was preserved in the untrained network. This indicates that the greater degree of invariance and equivariance for vertical reflection compared to 90° rotation is largely caused by the test images’ structure rather than task training. One interpretation is that, unlike 90°rotation, vertical and horizontal reflection both preserve the relative prevalence of vertical and horizontal edge energy, which may not be equal in natural images [29–32]. To test if the emergence of equivariance and invariance under horizontal reflection is unique to our controlled stimulus set (which contained many horizontally-symmetrical images), we repeated these analyses using natural images sampled from the ImageNet validation set (Fig. 3C-D). The training-dependent layer-by-layer increase in equivariance and invariance to horizontal reflection was as pronounced for natural images as it was for the rendered 3D object images. Therefore, the emergent invariance and equivariance under horizontal reflection are not an artifact of the synthetic object stimulus set.

Repeating these analyses on random noise images, the ImageNet-trained AlexNet still showed a slightly higher level of horizontal reflection-equivariance (Fig. 3E), demonstrating the properties of the features learned in the task independently of symmetry structure in the test images. When we evaluated an untrained AlexNet on random noise images (Fig. 3F), that is, when there was no structure in either the test stimuli or the network weights, the differences between horizontal reflection, vertical reflection, and rotation measures disappeared, and the invariance and equivariance measures were zero, as expected (see Fig. S7 for the distribution of equivariance and invariance across test images).

To summarize this set of analyses, a high level of reflection-invariance is associated with the layer’s pooling size and the reflection-equivariance of its feeding representation. These properties depend not only on the architecture but also on the network’s training. Training on recognizing objects in natural images induces a greater degree of invariance and equivariance to horizontal reflection compared to vertical reflection or 90° rotation. This is consistent with the statistics of natural images as experienced by an upright observer looking, along a horizontal axis, at upright bilaterally symmetric objects. Image reflection, in such a world ordered by gravity, does not change the category of an object (although rare examples of dependence of meaning on handedness exist, such as the letters p and q, and molecules whose properties depend on their chirality). However, the analyses reported thus far leave unclear whether natural image statistics alone or the need to disregard the handedness for categorization drive mirror-symmetric viewpoint tuning. In the following section, we examine what it is about the training that drives viewpoint tuning to be mirror-symmetric.

### Learning to discriminate among categories of bilaterally symmetric objects induces mirror-symmetric viewpoint tuning

To examine how task demand and visual diet influence mirror-symmetric viewpoint tuning, we trained four deep convolutional neural networks of the same architecture on different datasets and tasks (Fig. 4). The network architecture and training hyper-parameters are described in the Materials and Methods section (for training-related metrics, see Fig. S4). Once trained, each network was evaluated on the 3D object images used in Fig. 2, measuring mirror-symmetric viewpoint tuning qualitatively (Fig. 4B) and quantitatively (Fig. 4C). First, we considered a network trained on CIFAR-10 [33], a dataset of small images of 10 bilaterally symmetric categories (airplanes, cars, birds, cats, deer, dogs, frogs, horses, ships, and trucks). Although this dataset contains no human face images (such images appear coincidentally in the ImageNet dataset, [34]), the CIFAR-10-trained network reproduced the result of a considerable level of mirror-symmetric viewpoint tuning for faces in layers fc1 and fc2 (Fig. 4B, top row). This network also showed mirror-symmetric viewpoint tuning for other bilaterally symmetric objects such as cars, airplanes, and boats (Fig. 4C, blue lines).

**Figure 4.**
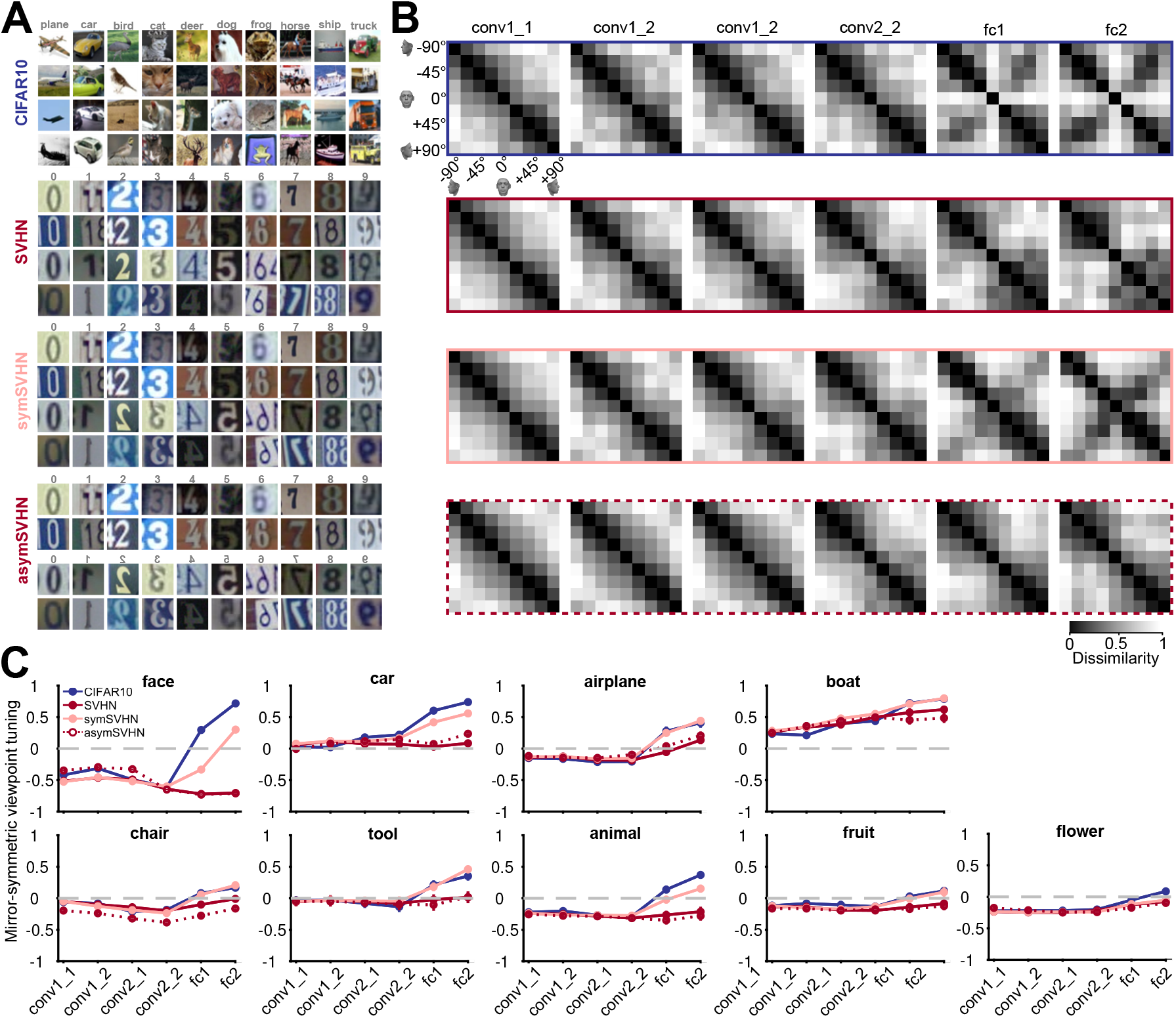
The effect of training task and training dataset on mirror-symmetric viewpoint tuning. (**A**) Four datasets are used to train deep neural networks of the same architecture: CIFAR-10, a natural image dataset with ten bilaterally symmetric object categories; SVHN, a dataset with mostly asymmetric categories (the ten numerical digits); symSVHN, a version of the SVHN dataset in which the categories were made bilaterally symmetric by horizontally reflecting half of the training images (so 7 and count as members of the same category); asymSVHN, the same image set as in symSVHN but with the mirrored images assigned to ten new distinct categories (so 7 and count as members of distinct categories). (**B**) Each row represents the RDMs of the face exemplar images from nine viewpoints for each trained network corresponding to its left side panel. Each entry of the RDM represents the dissimilarity (1 *−* Pearson’s r) between two pairs of image-induced activity vectors in the corresponding layer. The RDMs’ order from left to right refers to the depth of layers within the network. As the dissimilarity color bar indicates, the dissimilarity values increase from black to white color. (**C**) Mirror-symmetric viewpoint tuning index values across layers for nine object categories in each of the four networks. The solid circles refer to the average of the index across 25 exemplars within each object category for three networks trained on 10 labels. The red dashed line with open circles belongs to the asymSVHN network trained on 20 labels. The gray dashed lines indicate the index of zero. Error bars represent the standard error of the mean calculated across exemplars.

We then considered a network trained on SVHN (Street View House Numbers) [35], a dataset of photographs of numerical digits. Its categories are mostly asymmetric (since all ten digits except for ‘0’ and ‘8’ are asymmetric). Unlike the network trained on CIFAR-10, the SVHN-trained network showed a very low level of mirror-symmetric viewpoint tuning for faces. Furthermore, its levels of mirror-symmetric viewpoint tuning for cars, airplanes, and boats were reduced relative to the CIFAR-10-trained network.

SVHN differs from CIFAR-10 both in its artificial content and the asymmetry of its categories. To disentangle these two factors, we designed a modified dataset, “symSVHN”. Half of the images in symSVHN were horizontally reflected SVHN images. All of the images maintained their original category labels (e.g., images of ‘7’s and ‘ ‘s belonged to the same category). We found that the symSVHN-trained network reproduced the mirror-symmetric viewpoint tuning observed in the CIFAR-10-trained network.

Last, we modified the labels of symSVHN such that the flipped digits would count as 10 separate categories, in addition to the 10 unflipped digit categories. This dataset (“asymSVHN”) has the same images as symSVHN, but it is designed to require reflection-sensitive recognition. The asymSVHN-trained network reproduced the low levels of mirror-symmetric viewpoint tuning observed for the original SVHN dataset. Together, these results suggest that given the spatial pooling carried out by fc1, the task demand of *reflection-invariant recognition* is a sufficient condition for the emergence of mirror-symmetric viewpoint tuning for faces.

### Equivariant local features drive mirror-symmetric viewpoint tuning

What are the image-level visual features that drive the observed mirror-symmetric viewpoint tuning? Do mirror-reflected views of an object induce similar representations because of global 2D configurations shared between such views? Or alternatively, are reflection-equivariant local features sufficient to explain the finding of similar responses to reflected views in fc1?

We used a masking-based importance mapping technique [36] to characterize which features drive the responses of units with mirror-symmetric viewpoint tuning. First, we created importance maps whose elements represent how local features influence each unit’s response to different object views. The top rows of panels A and B in Fig. 5 show examples of such maps for two units, one that shows considerable mirror-symmetric viewpoint tuning for cars and another that shows considerable mirror-symmetric viewpoint tuning for faces. Next, we empirically tested whether the local features highlighted by the importance maps are sufficient and necessary for generating mirror-symmetric viewpoint tuning. We used two image manipulations: insertion and deletion [36] (Fig. 5A-B, middle rows). When we retained only the most salient pixels (i.e., insertion), we observed that the units’ mirror-symmetric viewpoint tuning levels were similar to those induced by unmodified images (Fig. 5A-B, dark blue lines). This result demonstrates that the local features suffice for driving mirror-symmetrically tuned responses. Conversely, greying out the most salient pixels (deletion) led to a complete loss of mirror-symmetric viewpoint tuning (Fig. 5A-B, red lines). This result demonstrates that the local features are necessary to drive mirror-symmetrically tuned responses. To examine this effect systematically, we selected one unit for each of the 225 3D objects that showed high mirror-symmetric viewpoint tuning. We then tested these 225 units with insertion and deletion images produced with different thresholds (Fig. 5C). Across all threshold levels, the response to insertion images was more similar to the response to unmodified images, whereas deletion images failed to induce mirror-symmetric viewpoint tuning.

**Figure 5.**
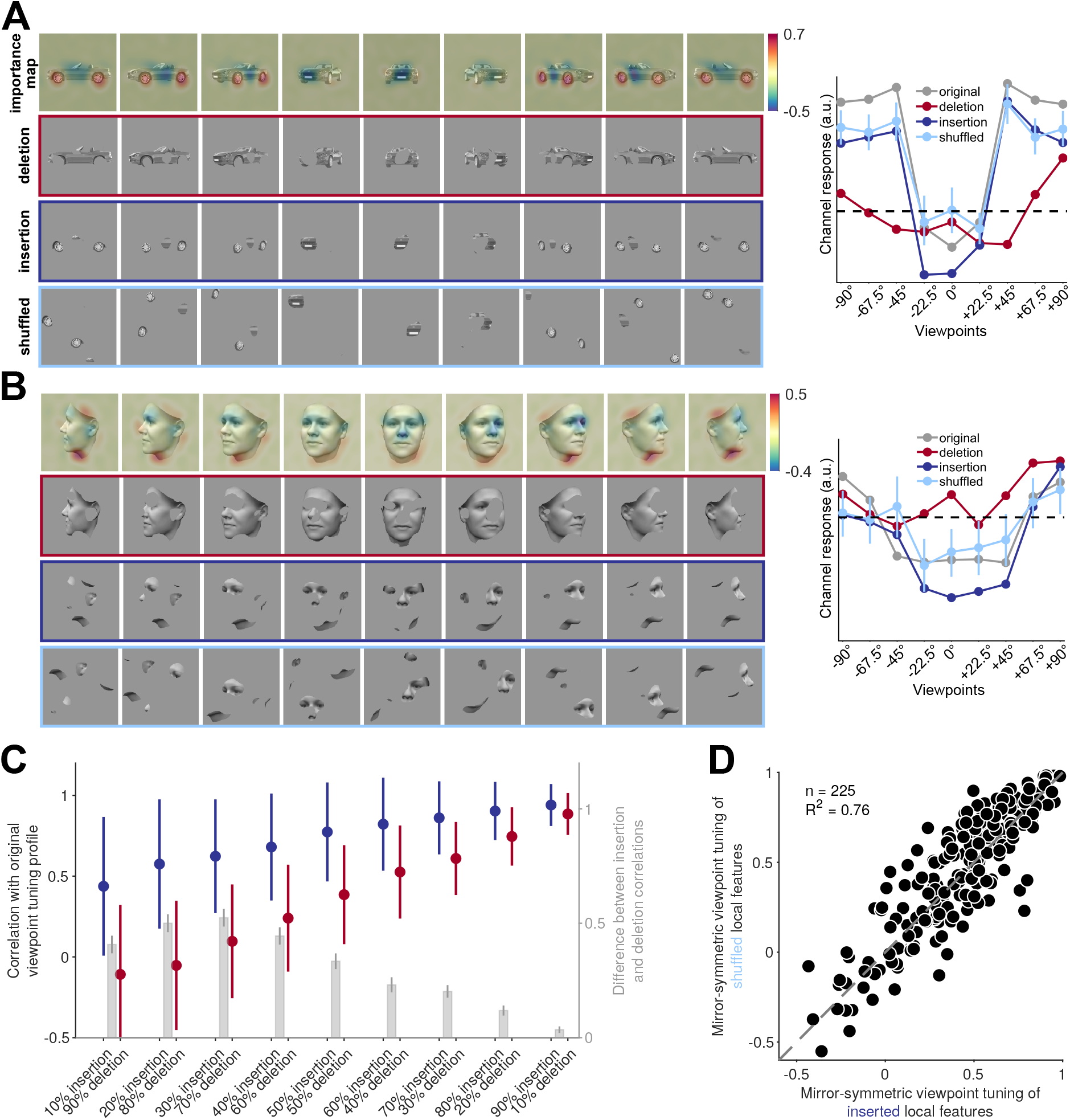
Reflection-invariant viewpoint-specific responses are driven mostly by local features. This figure traces image-level causes for the mirror-symmetric viewpoint tuning using Randomized Input Sampling for Explanation (RISE, [36]). (**A**) Analysis of the features of different views of a car exemplar that drive one particular unit in fully connected layer fc6 of AlexNet. The topmost row in each panel depicts an image-specific *importance map* overlaid to each view of the car, charting the contribution of each pixel to the unit’s response. The second row (“deletion”) depicts a version of each input image in which the 25 percent most contributing pixels are masked with the background gray color. The third row (“insertion”) depicts a version of the input images in which only the most contributing 25 percent of pixels appear. The last row represents the shuffled spatial configuration of extracted local features, which maintains their structure and changes their locations. The charts on the right depict the units’ responses to the original, deletion, insertion, and shuffled images. The dashed line indicates the units’ response to a blank image. (**B**) Analogous analysis of the features of different views of a face that drive a different unit in fully connected layer fc6 of AlexNet. (**C**) Testing local contributions to mirror-symmetric viewpoint tuning across all object exemplars and insertion/deletion thresholds. For each object exemplar, we selected a unit with a highly view-dependent but symmetric viewpoint tuning (the unit whose tuning function was maximally correlated with its reflection). We then measured the correlation between this tuning function and the tuning function induced by insertion or deletion images that were generated by a range of thresholding levels (from 10 to 90%). Note that each threshold level consists of images with the same number of non-masked pixels appearing in the insertion and deletion conditions. In the insertion condition, only the most salient pixels are retained, and in the deletion condition, only the least salient pixels are retained. The solid circles and error bars indicate the median and standard deviation over 225 objects, respectively. The right y-axis depicts the difference between insertion and deletion conditions. Error bars represent the SEM. (**D**) For each of 225 objects, we selected units with mirror-symmetric viewpoint tuning above the 95 percentile (*≈*200 units) and averaged their corresponding importance maps. Next, we extracted the top 25 percent most contributing pixels from the averaged maps (insertion) and shuffled their spatial configuration (shuffled). We then measured the viewpoint-RDMs for either the inserted or shuffled object image set. The scatterplot compares the mirror-symmetric viewpoint tuning index between insertion and shuffled conditions, calculated across the selected units. Each solid circle represents an exemplar object. The high explained variance indicates that the global configuration does not play a significant role in the emergence of mirror-symmetric viewpoint tuning.

These results indicate a role for local features in mirrorsymmetric tuning. However, the features may form larger-scale configurations synergistically. To test the potential role of such configurations, we shuffled contiguous pixel patches that were retained in the insertion condition. This manipulation destroyed global structure while preserving local features (Fig. 5A-B, bottom row). We found that the shuffled images largely preserved the units’ mirror-symmetric viewpoint tuning (Fig. 5D). Thus, it is the mere presence of a similar set of reflected local features (rather than a reflected global configuration) that explains most of the acquired mirror-symmetric viewpoint tuning. Note that such local features must be either symmetric at the image level (e.g., the wheel of a car at a side view), or induce a reflection-equivariant representation (e.g., an activation map that highlights profile views of a nose, regardless of their orientation). The fc6 layer learns highly symmetrical weight kernels, reducing the sensitivity to local feature configurations and enabling the generation of downstream reflection-invariant representations compared to convolutional layers (Fig. S5).

## Discussion

In this paper, we propose a simple learning-driven explanation for the mirror-symmetric viewpoint tuning for faces in the macaque AL face-patch. We found that CNNs trained on object recognition reproduce this tuning in their fully connected layers. Based on insilico experiments, we suggest two jointly sufficient conditions for the emergence of mirror-symmetric view-point tuning. First, training the network to discriminate among bilaterally symmetric objects yields reflection-equivariant representations in the deeper convolutional layers. Then, subsequent pooling of these reflection-equivariant responses by units with large receptive fields leads to reflection-invariant representations with mirror-symmetric view tuning similar to that observed in the AL face patch. Like our models, monkeys need to recognize bilaterally symmetric objects that are oriented by gravity with robustness to view, and the primate visual system can pool responses from earlier stages of representation to achieve this. We further show that in CNNs, such tuning is not limited to faces and occurs for multiple object categories with bilateral symmetry. This result yields a testable prediction for primate electrophysiology and fMRI.

### Mirror-symmetric viewpoint tuning in brains and machines

Several species, including humans, confuse lateral mirror images (e.g., the letters b and d) more often than vertical mirror images (e.g., the letters b and p) [37, 38]. Children often experience this confusion when learning to read and write [39– Single-cell recordings in macaque monkeys presented with simple stimuli indicate a certain degree of reflection-invariance in IT neurons [43, 44]. Human neuroimaging experiments also revealed reflection-invariance across higher-level visual regions for human heads [45–48] and other bilaterally symmetric objects [47, 49].

When a neuron’s response is reflection-invariant and yet the neuron responds differently to different object views, it is exhibiting mirror-symmetric viewpoint tuning. Such tuning has been reported in a small subset of monkeys’ STS and IT cells in early recordings [50, 51]. fMRI-guided single-cell recordings revealed the prevalence of this tuning profile among the cells of face patch AL [3]. The question of why mirror-symmetric viewpoint tuning emerges in the cortex has drawn both mechanistic and functional explanations. Mechanistic explanations suggest that mirror-symmetric viewpoint tuning is a by-product of increasing interhemispheric connectivity and receptive field sizes. Due to the anatomical symmetry of the nervous system and its cross-hemispheric interconnectivity, mirror-image pairs activate linked neurons in both hemispheres [52, 53]. A functional perspective explains partial invariance as a stepping stone toward achieving fully view-invariant object recognition [3]. Our results support a role for both of these explanations. We showed that global spatial pooling is a sufficient condition for the emergence of reflection-invariant responses, *if* the pooled representation is reflection-equivariant. Global average pooling extends the spatially integrated stimulus region. Likewise, interhemispheric connectivity may result in cells with larger receptive fields that cover both hemifields. We also showed that equivariance can be driven by the task demand of discriminating among objects that have bilateral symmetry. The combined effect of equivariance and pooling leads to a leap in reflection-invariance between the last convolutional layer and the fully connected layers in CNNs. This transition may be similar to the transition from view-selective cells in face patches ML/MF to mirror-symmetric viewpoint-selective cells in AL. In both CNNs and primate cortex, the mirror-symmetrically viewpoint-tuned neurons are a penultimate stage on the path to full view invariance [3].

### Unifying the computational explanations of mirror-symmetric viewpoint tuning

Two computational models have been suggested to explain AL’s mirror-symmetric viewpoint tuning, the first attributing it to Hebbian learning with Oja’s rule [18], the second to training a CNN to invert a face-generative model [13]. A certain extent of mirror-symmetric view-point tuning was also observed in CNNs trained on face identification (Figure 3E-ii in [13], Figure 2 in [11]). In light of the current work, these models can be viewed as special cases of a considerably more general class of models. Our results generalize the computational account in terms of the stimulus domain. Both [18] and [13] trained neural networks with face images. Here, we show that it is not necessary to train on a specific object category (including faces) in order to acquire reflection equivariance and invariance for exemplars of that category. Instead, learning mirror-invariant stimulus-to-response mappings gives rise to equivariant and invariant representations also for novel stimulus classes.

Our results also generalize the computational account of mirror-symmetric viewpoint tuning in terms of the model architecture. The two previous models incorporated the architectural property of spatial pooling: the inner product of inputs and synaptic weights in the penultimate layer of the HMAX-like model in [18] and the global spatial pooling in the f4 layer of the EIG model [13]. We showed that in addition to the task, such spatial pooling is an essential step toward the emergence of mirror-symmetric tuning in our findings.

### Limitations and future work

One limitation of the current study is the use of simpler, relatively shallow convolutional neural networks (such as AlexNet and VGG16, [21, 23]) rather than deeper ones (such as ResNet-50, [54]). We chose to focus on shallower networks since we required a simple macroscopic correspondence between the layer structure of the artificial networks and the ventral visual stream. In shallower CNNs such as AlexNet, the number of layers is similar to the number of feedforward steps in the visual cortex and the progression of representations across layers can be directly linked to the progression of representations across ventral stream regions [55]. While deeper networks with 50, 100 or even 200 layers can show higher quantitative correspondence with cortical representations [56, 57], this improved correspondence depends on allowing for arbitrary linear transformations between the neural network activations and the cortical representations. When neural networks and cortical responses are compared using more restrictive mappings such as non-weighted or weighted Representational Similarity Analysis, the deeper models lose their advantage as models of cortical representations [58]. For these reasons, we chose to focus on neural networks that can be compared to the ventral stream without additional mapping steps.

A second limitation of the current study is that we demonstrate the correspondence between the invariance properties of AL and fc1 activations only qualitatively. Future work, utilizing neuronal responses to multiple views of diverse visual objects, can quantitatively compare equivariance and invariance between biological neurons and neural network models. The analysis of such an experiment can take into account also the more powerful models that are formed by data-driven linear regression between neural network activations and neuronal responses.

Last, our findings are empirical in nature. Therefore, they might be limited to the particular design choices shared across the CNNs we evaluated. This limitation stands in contrast with the model proposed by Leibo and colleagues [18], which is reflection-invariant by construction. However, it is worth noting that the model proposed by Leibo and colleagues is reflection invariant only with respect to the horizontal center of the input image (Fig. S6). CNNs trained to discriminate among bilaterally symmetric categories develop mirror-symmetric viewpoint tuning across the visual field (Fig. S6). The latter result pattern is more consistent with the relatively position-invariant response properties of AL neurons (Fig. S10 in [3]).

### A novel prediction: mirror-symmetric viewpoint tuning for non-face objects

Mirror-symmetric viewpoint tuning has been mostly investigated using face images. Extrapolating from the results in CNNs, we hypothesize that mirror-symmetric viewpoint tuning for non-face objects should exist in cortical regions homologous to AL. The mirror-symmetric tuning of these objects does not necessarily have to be previously experienced by the animal.

This hypothesis is consistent with the recent findings of Bao and colleagues [59]. They report a functional clustering of IT into four separate networks. Each of these networks is elongated across the IT cortex and consists of three stages of processing. We hypothesize that the intermediate nodes of the three non-face selective networks have reflection-invariant yet view-selective tuning, homologous to AL’s representation of faces.

Our controlled stimulus set, which includes systematic 2D snapshots of 3D real-world naturalistic objects, is available online. Future electrophysiological and fMRI experiments utilizing this stimulus set can verify whether the mirror-symmetric viewpoint tuning for non-face categories we observe in task-trained CNNs also occurs in the primate IT.

## Methods

### 3D object stimulus set

We generated a diverse image set of 3D objects rendered from multiple views in the depth rotation. Human faces were generated using the Basel Face Model [60]. For the non-face objects, we purchased access to 3D models on TurboSquid (http://www.turbosquid.com). The combined object set consisted of nine categories (cars, boats, faces, chairs, airplanes, animals, tools, fruits, and flowers). Each category included 25 exemplars. We rendered each exemplar from nine views, giving rise a total of 2,025 images. The views span from -90° (left profile) to +90°, with steps of 22.5°. The rendered images were converted to grayscale, placed on a uniform gray background, and scaled to 227 ×227 pixels to match the input image size of AlexNet, or to 224 ×224 to match the input image size of the VGG-like network architectures. Mean luminance and contrast of non-background pixels were equalized across images using the SHINE toolbox [61].

### Pre-trained neural networks

We evaluated an ImageNet-trained AlexNet [21], an ImageNet-trained VGG16 [23], and VGGFace–a similar architecture to VGG16, trained on the VGG Face dataset [24]. We chose these networks based on their simple architectures and the rough correspondence between their layer counts and the number of feedforward steps in the visual cortex. The architecture of VGG16 is deeper than AlexNet. Both were trained on the ImageNet dataset [62], which consists of ∼1.2 million natural images from 1000 object categories (available on Matlab Deep Learning Toolbox and Pytorch frameworks, [63, 64]). The VGGFace model was trained on ∼2.6 million face images from 2622 identities (available on the MatConvNet library, [65]). Depending on the architecture, each network has a different number of convolutional (conv), max-pooling (pool), rectified linear unit (relu), normalization (norm), and fully connected (fc) layers. For untrained networks, we initialized the weights and biases using a random Gaussian distribution with a zero mean and a variance inversely proportional to the number of inputs per unit [66].

### Trained-from-scratch neural networks

To control for the effects of the training task and “visual diet”, we trained four networks employing the same convolutional architecture on four different datasets: CIFAR-10, SVHN, symSVHN, and asymSVHN.

#### CIFAR-10

CIFAR-10 consists of 60,000 RGB images of 10 classes (airplane, automobile, bird, cat, deer, dog, frog, horse, ship, truck) downscaled to 32 ×32 pixels [33]. We randomly split CIFAR-10’s designated training set into 45,000 images used for training and 5,000 images used for validation. No data augmentation was employed. The reported classification accuracy (Fig. S4) was evaluated on the remaining 10,000 CIFAR-10 test images.

#### SVHN

SVHN [35] contains 99,289 RGB images of 10 digits (0 to 9) taken from real-world house number photographs [35], cropped to character bounding boxes and downsized to 32 ×32 pixels. We split the dataset into 73,257 images for the training set and 26,032 images for the test set. As with the CIFAR-10 dataset, we randomly selected 10 percent of training images as the validation set.

#### symSVHN and asymSVHN

As a control experiment, we horizontally flipped half of the SVHN training images while keeping their labels unchanged. This manipulation encouraged the model trained on these images to become reflection-invariant in its decisions. This dataset was labeled as “symSVHN”.

In a converse manipulation, we applied the same horizontal flipping but set the flipped images’ labels to ten new classes. Therefore, each image in this dataset pertained to one of 20 classes. This manipulation removed the shared response mapping of mirror-reflected images and encouraged the model trained on these images to become sensitive to the reflection operation. This dataset was labeled as “asymSVHN”.

#### Common architecture and training procedure

The networks’ architecture resembled the VGG architecture. It contained two convolutional layers followed by a maxpooling layer, two additional convolutional layers, and three fully connected layers. The size of convolutional filters was set to 3 ×3 with a stride of 1. The four convolutional layers consisted of 32, 32, 64, and 128 filters, respectively. The size of the max-pooling window was set to 2 ×2 with a stride of 2. The fully-connected layers had 128, 256, and 10 channels and were followed by a softmax operation (the asymSVHN network had 20 channels in its last fully connected layer instead of 10). We added a batch normalization layer after the first and the third convolutional layers and a dropout layer (probability = 0.5) after each fully-connected layer to promote quick convergence and avoid overfitting.

The networks’ weights and biases were initialized randomly using the uniform He initialization [67]. We trained the models using stochastic gradient descent (SGD). The CIFAR-10 network was trained for 250 epochs with a learning rate of 2.5 ×10^*−*4^, a momentum of 0.9, a batch size of 128 images, and a weight decay of 10^*−*4^. The SVHN/symSVHN/asymSVHN networks were trained for 200 epochs with a learning rate of 10^*−*4^, a momentum of 0.9, a batch size of 64 images, and a weight decay of 5 ×10^*−*5^. The hyper-parameters were determined using the validation data. The models reached around 85% test accuracy (CIFAR-10: 81%, SVHN: 92%, symSVHN: 88%, asymSVHN: 80%). Fig. S4 shows the models’ learning curves.

### Measuring representational dissimilarities

For the analyses described in Figures 2, 3, and 4, we first normalized the activation level of each individual neural network unit by subtracting its mean response level across all images of the evaluated dataset and dividing it by its standard deviation. The dissimilarity between the representations of two stimuli in a particular neural network layer (Figs. 2 and 4) was quantified as one minus the Pearson linear correlation coefficient calculated across all of the layer’s units (i.e., across the flattened normalized activation vectors). The *similarity* between representations (Fig. 3) was quantified by the linear correlation coefficient itself.

### Measuring mirror-symmetric viewpoint tuning

Using the representational dissimilarity measure described above, we generated an *n* × *n* dissimilarity matrix for each exemplar object *i* and layer *ℓ*, where *n* is the number of views (9 in our dataset). Each element of the matrix, 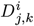, denotes the representational distance between views *j* and *k* of object exemplar *i*. The views are ordered such that *j* and *n* + 1 − *k* refer to horizontally reflected views.

We measured the mirror-symmetric viewpoint tuning index of the resulting RDMs by

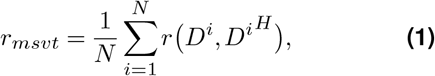

where *r* (·,·) is the Pearson linear correlation coefficient across view pairs, *D*^*H*^ refers to horizontally flipped matrix such that 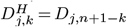, and *N* refers to number of object exemplars. The frontal view (which is unaltered by reflection) was excluded from this measure to avoid spurious inflation of the correlation coefficient.

### Measuring equivariance and invariance

Representational equivariance and invariance were measured for an ImageNet-trained AlexNet and an untrained AlexNet with respect to three datasets: the 3D object image dataset described above, a random sample of 2,025 ImageNet test images, and a sample of 2,025 random noise images (Fig. 3). Separately for each layer *ℓ* and image set *x*_1_, …, *x*_2025_, we measured invariance by

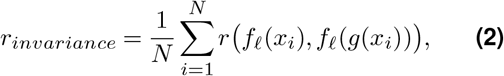

where *f*_*ℓ*_(·) is the mapping from an input image *x* to unit activations in layer *lℓ, g*(·) is the image transformation of interest–vertical reflection, horizontal reflection, or rotation and *r* is the Pearson linear correlation coefficient calculated across units, flattening the units’ normalized activations into a vector in the case of convolutional layers.

In order to estimate equivariance, we used the following definition:

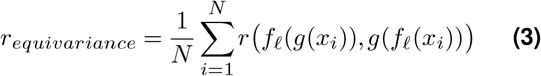

Note that in this case, *g*(·) was applied both to the input images and the feature maps. This measure can be viewed as the inverse of an additive realization of latent space G-empirical equivariance deviation (G-EED) [25]. To prevent spurious correlations that may result from flipping and rotating operations, we have removed the central column when flipping horizontally, the central row when flipping vertically, and the central pixel when rotating 90 degrees. As a result, any correlations we observe are unbiased.

### Importance mapping

We used an established masking-based importance mapping procedure [36] to identify visual features that drive units that exhibit mirror-symmetric viewpoint tuning profiles. Given an object for which the target unit showed mirror-symmetric viewpoint tuning, we dimmed the intensities of the images’ pixels in random combinations to estimate the importance of image features. Specifically, for each image, we generated 5000 random binary masks. Multiplying the image with these masks yielded 5000 images in which different subsets of pixels were grayed out. These images were then fed to the network as inputs. The resulting importance maps are averages of these masks, weighted by target unit activity. To evaluate the explanatory power of the importance map of each stimulus, we sorted the pixels according to their absolute values in the importance map and identified the top quartile of salient pixels. We then either retained (“insertion”) or grayed out (“deletion”) these pixels, and the resulting stimulus was fed into the network (Fig. 5A-B). Due to the uniform gray background, we only considered foreground pixels. A second analysis compared viewpoint tuning between original images, deletion images, and insertion images across 10 thresholds, from 10% to 90%, with steps of 10% (Fig. 5C).

We conducted an additional analysis to examine the influence of global structure on the mirror-symmetric viewpoint tuning of the first fully connected layer (Fig. 5D). To conduct this analysis at the unit population level, we generated one insertion image-set per object. First, we correlated each unit’s view tuning curve against a V-shaped tuning template (i.e., a response proportional to the absolute angle of deviation from a frontal view) and retained only the units with positive correlations. We then correlated each unit’s view-tuning curve with its reflected counterpart. We selected the top 5% most mirror-symmetric units (i.e., those showing the highest correlation coefficients).

For each object view, we generated an importance map for each of the selected units and averaged these maps across units. Using this average importance map, we generated an insertion image by retaining the top 25% most salient pixels. To test the role of global configuration, we generated a shuffled version of each insertion image by randomly relocating connected components. To assess model response to these images for each object exemplar, we computed the corresponding (9 ×9 views) RDM of fc1 responses given either the insertion images or their shuffled versions and quantified the mirror-symmetric viewpoint tuning of each RDM.

## ACKNOWLEDGEMENTS

Research reported in this publication was supported by the National Eye Institute of the National Institutes of Health under Award Numbers R01EY021594 and R01EY029998; by the National Institute Of Neurological Disorders And Stroke of the National Institutes of Health under Award Number RF1NS128897; and by the Department of the Navy, Office of Naval Research under ONR award number N00014-20-1-2292. This publication was made possible in part with the support of the Charles H. Revson Foundation to TG. The content is solely the responsibility of the authors and does not necessarily represent the official views of the National Institutes of Health or the Charles H. Revson Foundation. We acknowledge Dr. T. Vetter, Department of Computer Science, and the University of Basel, for the Basel Face Model.

## AUTHOR CONTRIBUTIONS

AF, WZ, WF, NK, and TG designed the research. AF performed the research. AF, NK, and TG wrote the paper. All authors have edited and approved the manuscript for submission.

## DATA AND CODE AVAILABILITY

The stimulus set and the source code required for reproducing our results will be available at the following link: https://github.com/amirfarzmahdi/AL-Symmetry.

## Supplementary Information

**Figure S1.**
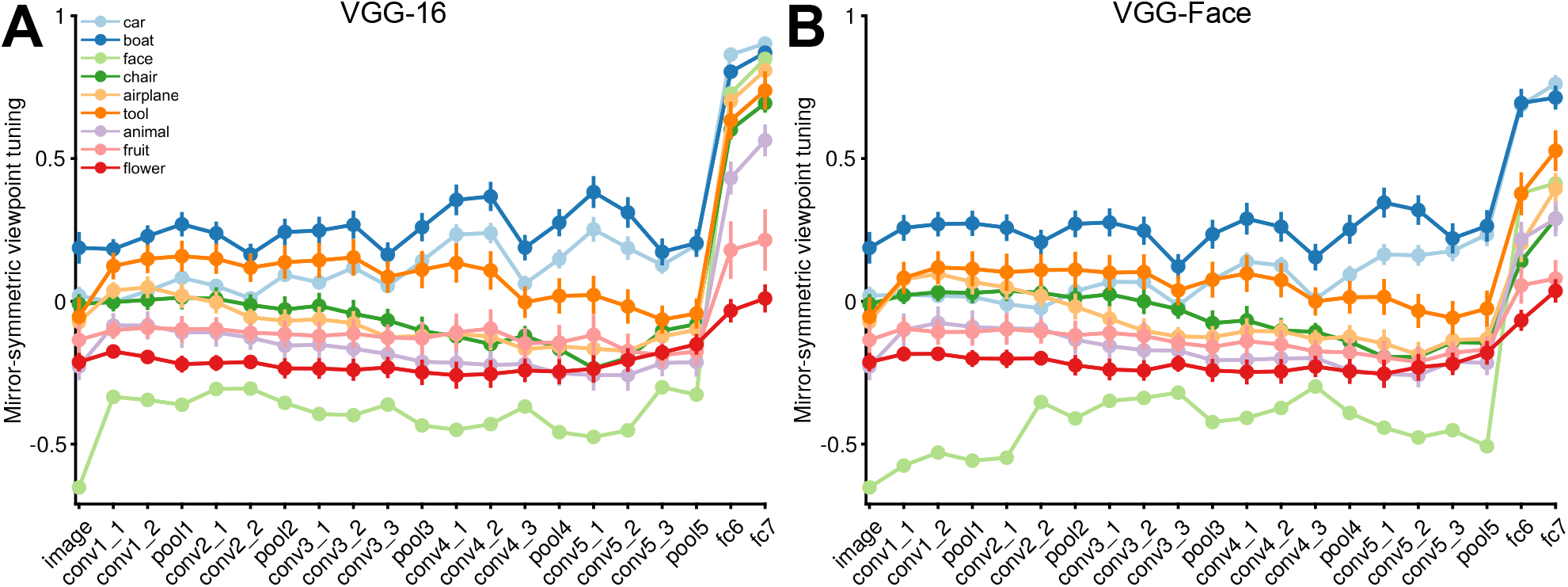
Similar layer-wise mirror-symmetric viewpoint tuning profiles are observed in networks of different architectures and training objectives. (**A**) The colored curves depict the mirror-symmetric viewpoint tuning indices across nine object categories (car, boat, face, chair, airplane, animal, tool, fruit, flower) in the VGG-16 layers. Each solid circle refers to the average of the index over 25 exemplars within each object category. Error bars represent the standard error of the mean. (**B**) Same analysis as (**A**), but for the Parkhi 2015 “VGG-Face” network [24].

**Figure S2.**
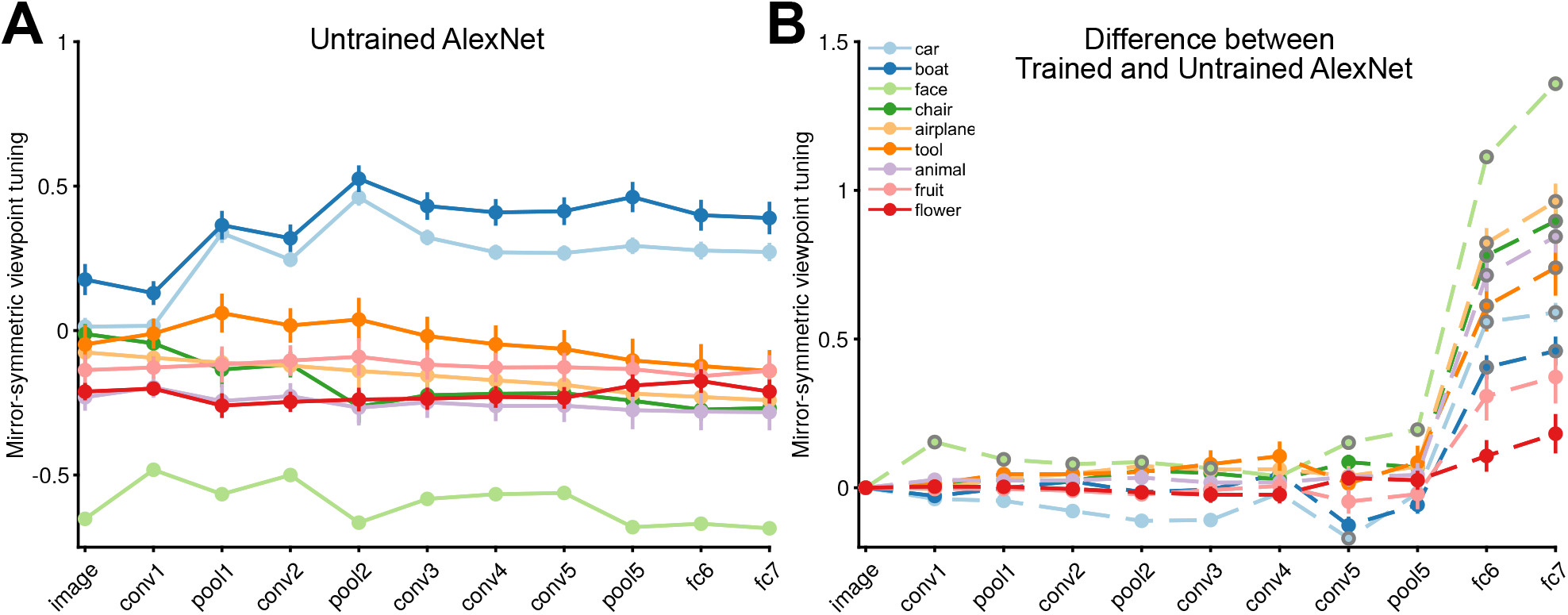
The mirror-symmetric viewpoint tuning index remains unchanged as the signal moves into the fully connected layers of the untrained network. (**A**) Each solid circle represents the average index for 25 exemplars within each object category (car, boat, face, chair, airplane, animal, tool, fruit, flower) for the untrained AlexNet network. (**B**) Each solid circle refers to the difference between the mirror-symmetric viewpoint tuning index of the trained versus the untrained AlexNet network. We evaluated the difference using the rank-sum test. We used the False discovery rate (FDR) correction for controlling 90 comparisons at q < .05 (9 categories and 10 layers, excluding the input layer, as it is the same in both networks). The solid circles with gray outlines indicate where the difference after FDR adjustment is significant. Error bars indicate the standard error of the mean.

**Figure S3.**
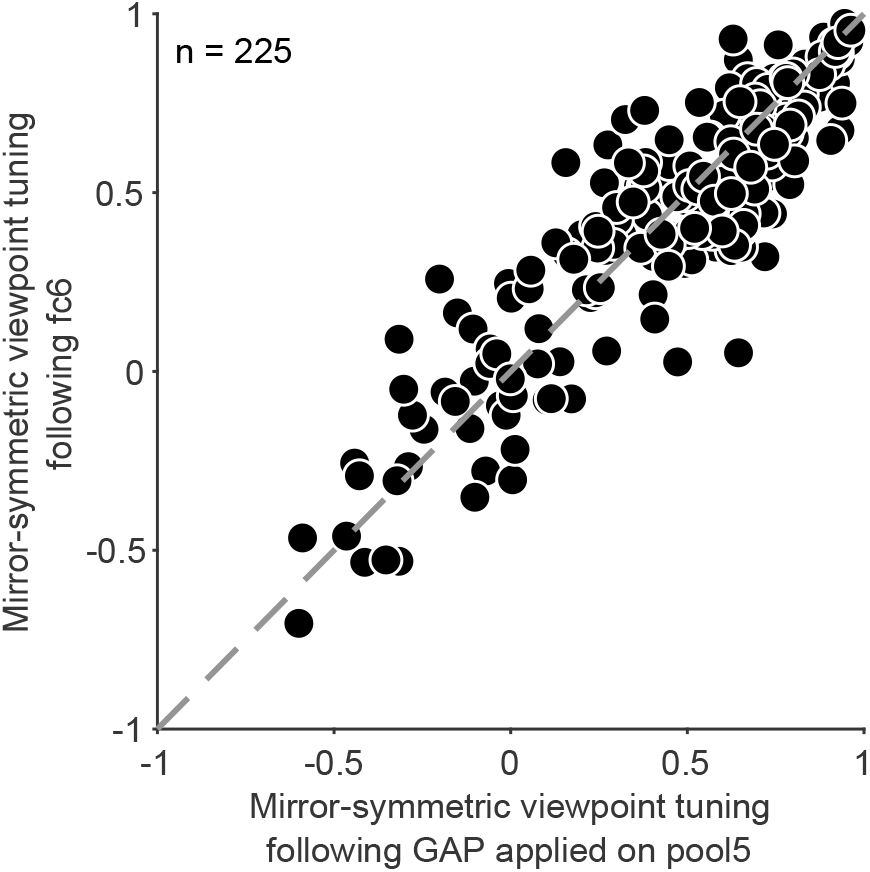
One of the key operations in fully-connected layers is spatial pooling. We analyzed the impact of this operation by artificially introducing global average pooling (GAP) instead of the first fully-connected layer (fc6) of ImageNet-trained AlexNet. Each element of the GAP representation refers to a spatial average of unit activations of one pool5 feature map. The scatterplot shows the mirror-symmetric viewpoint tuning index of GAP applied to pool5 (x-axis) relative to an fc6 representation (y-axis). Each circle represents one exemplar object. These results indicate that global spatial pooling introduced instead of fc6 is sufficient for rendering the pool5 representation mirror-symmetric viewpoint selective, reproducing the symmetry levels of the different fc6 view tuning curves across objects.

**Figure S4.**
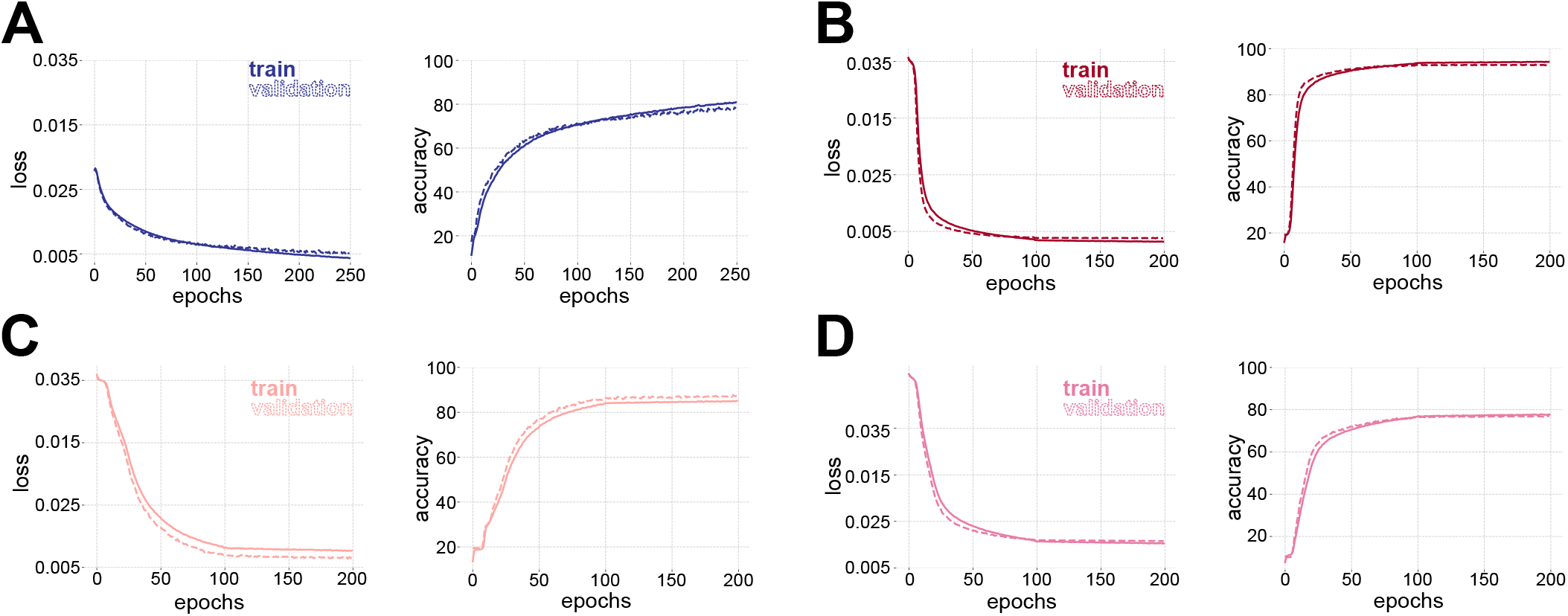
Network learning curves. (**A-D**) The loss and accuracy curves for the networks trained by CIFAR-10 (A), SVHN (B), symSVHN (C), asymSVHN (D) datasets. The x-axis denotes training epochs.

**Figure S5.**
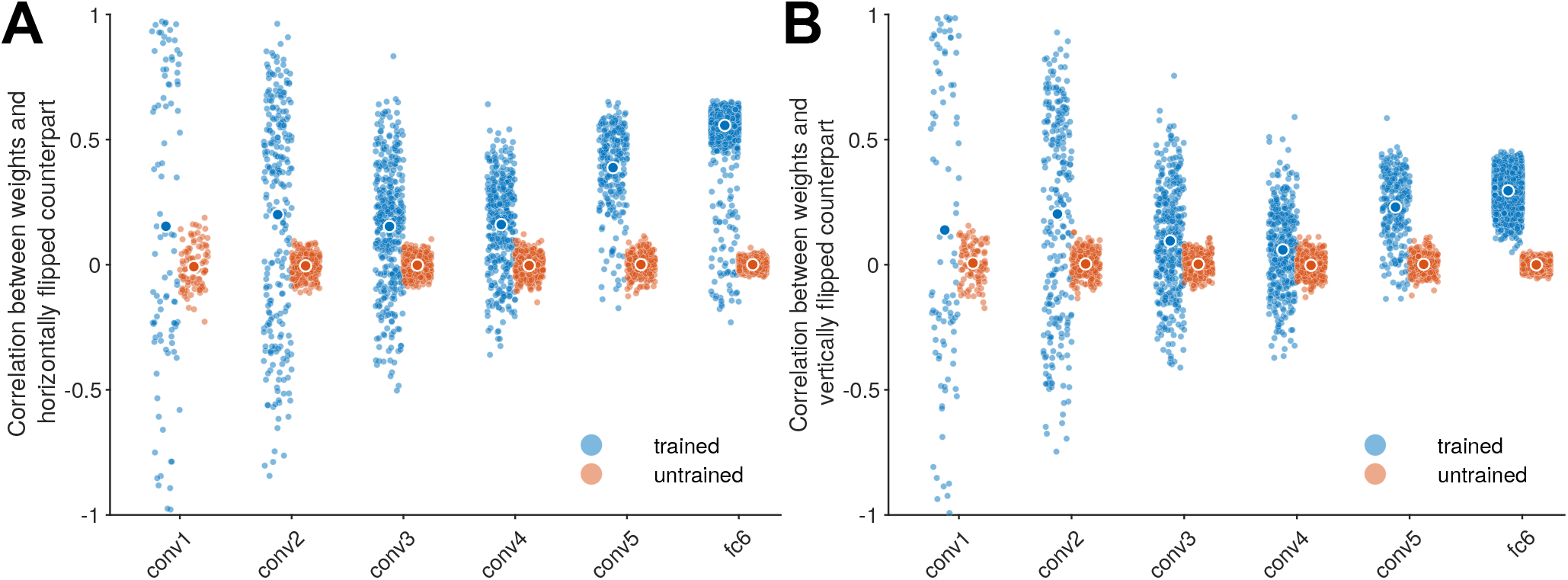
The emergence of mirror symmetric weight tensors in AlexNet. In order to examine the symmetry of neural network weights, we measured the linear correlation between each convolutional weight kernel and its horizontally and vertically flipped counterpart. To avoid replicated observations in the correlation analysis, we considered only the left (or top) half of the matrix, and excluded the central column (or row). Each dot represents one channel. This measurement was done for each convolutional layer in an AlexNet trained on ImageNet, as well as in an untrained AlexNet. The symmetry of the incoming weights to fc6 was evaluated in a similar fashion (note that the weights leading into this layer still have an explicit spatial layout, unlike fc7 and fc8). This analysis demonstrates that in the ImageNet-trained AlexNet network, weight symmetry increases with depth. Note that ImageNet training induces some highly asymmetrical kernels in conv1 and conv2. Together, these results suggest that while asymmetrical filters are useful low-level representations, the trained network incorporates symmetric weight kernels to generate downstream reflection-invariant representations.

**Figure S6.**
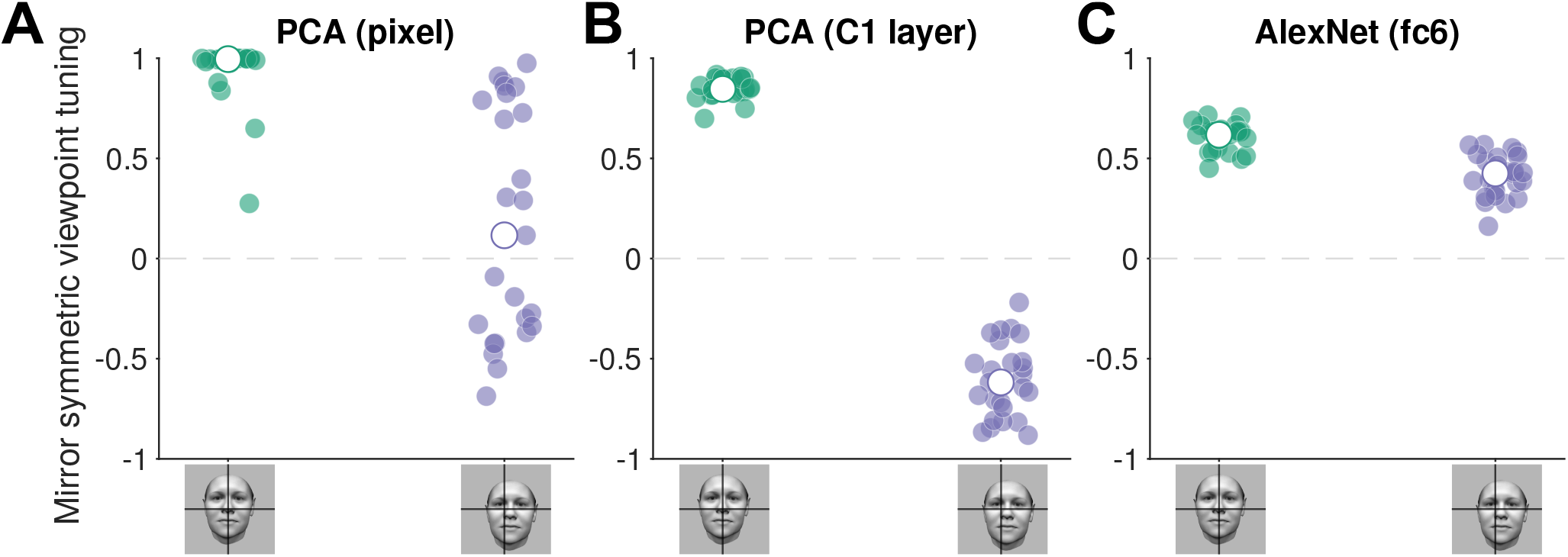
Comparison of mirror-symmetric viewpoint tuning in a supervised, PCA-based model [18] and a supervised CNN (AlexNet) trained on object recognition. Panels A and B depict how mirror-symmetric viewpoint tuning in a re-implementation of the Leibo and colleagues model [18] sharply declines for off-center test stimuli. In contrast, the same shift in center of the test stimuli has only a negligible effect on mirror-symmetric viewpoint tuning in the AlexNet trained on object recognition (Panel C). Implementation details: To reproduce the model described in [18], we generated a training stimulus set using the Basel Face Model. The stimulus set consisted of untextured synthetic faces of 40 identities, each depicted from 39 viewpoints. For panel A, we estimated a PCA of the pixel-space representation of this stimulus set. For panel B, we estimated a PCA of the stimulus set’s HMAX C1 layer representation. In both cases, the resulting latent representation had 1560 features (40*×*39). To test the model, we used the face stimulus set containing 25 exemplars in 9 viewpoints employed in Fig. 2. The viewpoints ranged from -90 to 90 degrees, with a step of 22.5°. Mirror-symmetric viewpoint tuning was extracted from a representational dissimilarity matrix (RDM) created per exemplar. Green and purple circles represent mirror-symmetric viewpoint tuning in centered and shifted images (with 15-pixel shifts in the x and y axes), respectively. White circles indicate the mean across all exemplars.

**Figure S7.**
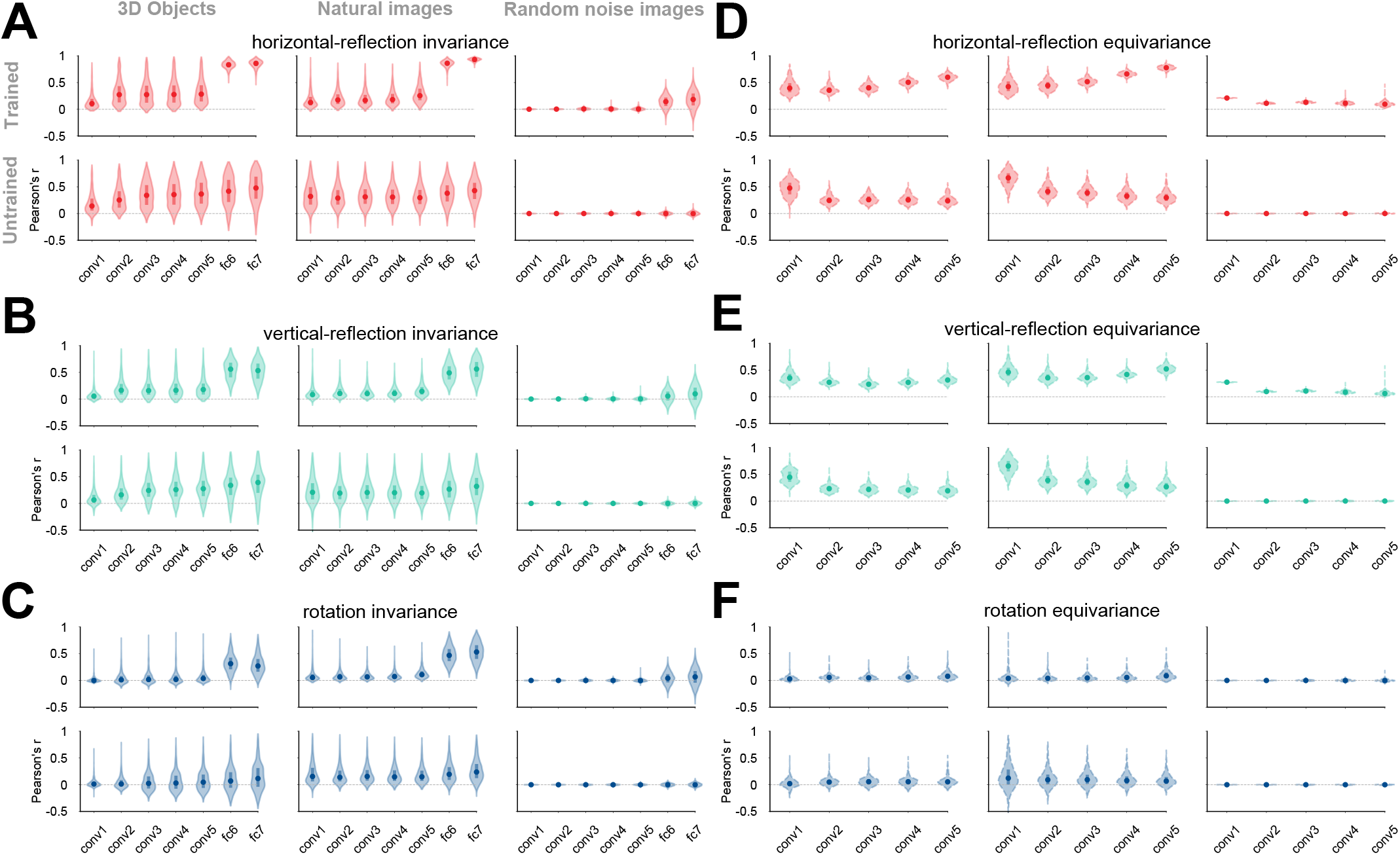
The distribution of equivariance and invariance across test images (3D object renders, natural images, or random noise images) in the representations of a deep convolutional neural network (AlexNet) trained on ImageNet or left untrained. Each violin plot depicts the distribution of invariance (panels A-C) or equivariance (D-F) across 2025 images. The different hues denote different transformations against which the equivariance and invariance were measured: horizontal flipping (red), vertical flipping (green), or 90°rotation (blue). The solid circles denote the median and the thick bars, the first and third quantiles. Invariance is a measure of the similarity between the activity pattern elicited by an image and the activity pattern elicited by a transformed version of the image. Equivariance is a measure of the similarity between the activity pattern elicited by a transformed image and a transformed version of the activity pattern of the untransformed image. Panels A, B, and C show the invariance over horizontally flipped, vertically flipped, and 90°rotated images, respectively. Panels D, E, and F depict the equivariance over the same transformations. **ImageNet training induces equivariance (in convolutional layers) and invariance (in fully connected layers) to the horizontal reflection of most natural images. This effect is less pronounced for vertical reflection and 90° rotation**.

## Bibliography

1. Doris Y Tsao, Winrich A Freiwald, Roger BH Tootell, and Margaret S Livingstone. A cortical region consisting entirely of face-selective cells. Science, 311(5761):670–674, 2006. doi:10.1126/science.1119983.

2. Sebastian Moeller, Winrich A Freiwald, and Doris Y Tsao. Patches with links: a unified system for processing faces in the macaque temporal lobe. Science, 320(5881):1355–1359, 2008. doi:10.1126/science.1157436.

3. Winrich A Freiwald and Doris Y Tsao. Functional compartmentalization and viewpoint generalization within the macaque face-processing system. Science, 330(6005):845–851, 2010. doi:10.1126/science.1194908.

4. Janis K Hesse and Doris Y Tsao. The macaque face patch system: a turtle’s underbelly for the brain. Nature Reviews Neuroscience, 21(12):695–716, 2020. doi:10.1038/s41583-020-00393-w.

5. Winrich A Freiwald. The neural mechanisms of face processing: cells, areas, networks, and models. Current Opinion in Neurobiology, 60:184–191, 2020. doi:10.1016/j.conb.2019.12.007.

6. Elias B Issa and James J DiCarlo. Precedence of the eye region in neural processing of faces. Journal of Neuroscience, 32(47):16666–16682, 2012. doi:10.1523/JNEUROSCI.2391-12.2012.

7. Daniel L. K. Yamins, Ha Hong, Charles F. Cadieu, Ethan A. Solomon, Darren Seibert, and James J. DiCarlo. Performance-optimized hierarchical models predict neural responses in higher visual cortex. Proceedings of the National Academy of Sciences, 111(23):8619–8624, 2014. doi:10.1073/pnas.1403112111.

8. Seyed-Mahdi Khaligh-Razavi and Nikolaus Kriegeskorte. Deep supervised, but not unsupervised, models may explain it cortical representation. PLOS Computational Biology, 10(11):1–29, 11 2014. doi:10.1371/journal.pcbi.1003915.

9. Amirhossein Farzmahdi, Karim Rajaei, Masoud Ghodrati, Reza Ebrahimpour, and Seyed-Mahdi Khaligh-Razavi. A specialized face-processing model inspired by the organization of monkey face patches explains several face-specific phenomena observed in humans. Scientific reports, 6 (1):1–17, 2016. doi:10.1038/srep25025.

10. Naphtali Abudarham, Idan Grosbard, and Galit Yovel. Face recognition depends on specialized mechanisms tuned to view-invariant facial features: Insights from deep neural networks optimized for face or object recognition. Cognitive Science, 45(9):e13031, 2021. doi:10.1111/cogs.13031.

11. Rajani Raman and Haruo Hosoya. Convolutional neural networks explain tuning properties of anterior, but not middle, face-processing areas in macaque inferotemporal cortex. Communications Biology, 3(1):221, May 2020. ISSN 2399-3642. doi:10.1038/s42003-020-0945-x. URL https://www.nature.com/articles/s42003-020-0945-x.

12. Le Chang, Bernhard Egger, Thomas Vetter, and Doris Y. Tsao. Explaining face representation in the primate brain using different computational models. Current Biology, 31(13):2785–2795.e4, 2021. doi:10.1016/j.cub.2021.04.014.

13. Ilker Yildirim, Mario Belledonne, Winrich Freiwald, and Josh Tenenbaum. Efficient inverse graphics in biological face processing. Science Advances, 6(10):eaax5979, 2020. doi:10.1126/sciadv.aax5979.

14. Haruo Hosoya and Aapo Hyvärinen. A mixture of sparse coding models explaining properties of face neurons related to holistic and parts-based processing. PLOS Computational Biology, 13(7):1–27, July 2017. doi:10.1371/journal.pcbi.1005667. Publisher: Public Library of Science.

15. Irina Higgins, L. Chang Victoria Langston, Demis Hassabis, Christopher Summerfield, Doris Tsao, and Matthew Botvinick. Unsupervised deep learning identifies semantic disentanglement in single inferotemporal face patch neurons. Nature Communications, 12(1):6456, December 2021. doi:10.1038/s41467-021-26751-5.

16. Seungdae Baek, Min Song, Jaeson Jang, Gwangsu Kim, and Se-Bum Paik. Face detection in untrained deep neural networks. Nature Communications, 12(1):7328, December 2021. doi:10.1038/s41467-021-27606-9.

17. Katharina Dobs, Julio Martinez, Alexander J. E. Kell, and Nancy Kanwisher. Brain-like functional specialization emerges spontaneously in deep neural networks. Science Advances, 8(11):eabl8913, 2022. doi:10.1126/sciadv.abl8913.

18. Joel Z Leibo, Qianli Liao, Fabio Anselmi, Winrich A Freiwald, and Tomaso Poggio. View-tolerant face recognition and hebbian learning imply mirror-symmetric neural tuning to head orientation. Current Biology, 27(1):62–67, 2017. doi:10.1016/j.cub.2016.10.015.

19. Maximilian Riesenhuber and Tomaso Poggio. Hierarchical models of object recognition in cortex. Nature Neuroscience, 2(11):1019–1025, November 1999. doi:10.1038/14819.

20. Erkki Oja. Simplified neuron model as a principal component analyzer. Journal of mathematical biology, 15(3):267–273, 1982. doi:10.1007/BF00275687.

21. Alex Krizhevsky, Ilya Sutskever, and Geoffrey E Hinton. ImageNet classification with deep convolutional neural networks. In Advances in Neural Information Processing Systems 25, pages 1097–1105. Curran Associates, Inc., 2012. doi:10.1145/3065386.

22. Nikolaus Kriegeskorte, Marieke Mur, and Peter Bandettini. Representational similarity analysis - connecting the branches of systems neuroscience. Frontiers in Systems Neuroscience, 2:4, 2008. doi:10.3389/neuro.06.004.2008.

23. Karen Simonyan and Andrew Zisserman. Very deep convolutional networks for large-scale image recognition. In International Conference on Learning Representations, 2015. doi:10.48550/arXiv.1409.1556.

24. Omkar M. Parkhi, Andrea Vedaldi, and Andrew Zisserman. Deep face recognition. In Proceedings of the British Machine Vision Conference (BMVC), pages 41.1–41.12. BMVA Press, September 2015. doi:10.5244/C.29.41.

25. Henry Kvinge, Tegan Emerson, Grayson Jorgenson, Scott Vasquez, Timothy Doster, and Jesse Lew. In what ways are deep neural networks invariant and how should we measure this? In Alice H. Oh, Alekh Agarwal, Danielle Belgrave, and Kyunghyun Cho, editors, Advances in Neural Information Processing Systems, 2022. URL https://openreview.net/forum?id=SCD0hn3kMHw.

26. Aharon Azulay and Yair Weiss. Why do deep convolutional networks generalize so poorly to small image transformations? Journal of Machine Learning Research, 20(184):1–25, 2019. doi:10.48550/arXiv.1805.12177.

27. Taco Cohen and Max Welling. Group equivariant convolutional networks. In International conference on machine learning, pages 2990–2999. PMLR, 2016. URL http://proceedings.mlr.press/v48/cohenc16.html.

28. Maurice Weiler, Fred A Hamprecht, and Martin Storath. Learning steerable filters for rotation equivariant cnns. In Proceedings of the IEEE Conference on Computer Vision and Pattern Recognition, pages 849–858, 2018. URL https://openaccess.thecvf.com/content_cvpr_2018/html/Weiler_Learning_Steerable_Filters_CVPR_2018_paper.html.

29. David M. Coppola, Harriett R. Purves, Allison N. McCoy, and Dale Purves. The distribution of oriented contours in the real world. Proceedings of the National Academy of Sciences, 95(7):4002–4006, 1998. doi:10.1073/pnas.95.7.4002.

30. Antonio Torralba and Aude Oliva. Statistics of natural image categories. Network: Computation in Neural Systems, 14(3):391–412, jan 2003. doi:10.1088/0954-898x_14_3_302.

31. Margaret Henderson and John T. Serences. Biased orientation representations can be explained by experience with nonuniform training set statistics. Journal of Vision, 21(8):10–10, 08 2021. doi:10.1167/jov.21.8.10.

32. Ahna R Girshick, Michael S Landy, and Eero P Simoncelli. Cardinal rules: visual orientation perception reflects knowledge of environmental statistics. Nature Neuroscience, 14(7):926–932, July 2011. doi:10.1038/nn.2831.

33. Alex Krizhevsky and Geoffrey Hinton. Learning multiple layers of features from tiny images. Technical Report 0, University of Toronto, Toronto, Ontario, 2009. URL https://www.cs.toronto.edu/~kriz/learning-features-2009-TR.pdf.

34. Kaiyu Yang, Jacqueline H. Yau, Li Fei-Fei, Jia Deng, and Olga Russakovsky. A study of face obfuscation in ImageNet. In Proceedings of the 39th International Conference on Machine Learning, volume 162 of Proceedings of Machine Learning Research, pages 25313–25330. PMLR, 17–23 Jul 2022. URL https://proceedings.mlr.press/v162/yang22q.html.

35. Yuval Netzer, Tao Wang, Adam Coates, Alessandro Bissacco, Bo Wu, and Andrew Y. Ng. Reading digits in natural images with unsupervised feature learning. In NIPS Workshop on Deep Learning and Unsupervised Feature Learning 2011, 2011. URL https://research.google/pubs/pub37648/.

36. Vitali Petsiuk, Abir Das, and Kate Saenko. Rise: Randomized input sampling for explanation of black-box models. In British Machine Vision Conference (BMVC), 2018. doi:10.48550/arXiv.1806.07421.

37. NS Sutherland. Visual discrimination of orientation by octopus: Mirror images. British Journal of Psychology, 51(1):9–18, 1960. doi:10.1111/j.2044-8295.1960.tb00719.x.

38. David C Todrin and Donald S Blough. The discrimination of mirror-image forms by pigeons. Perception & Psychophysics, 34(4):397–402, 1983. doi:10.3758/BF03203053.

39. Rosemery O Nelson and Arthur Peoples. A stimulus-response analysis of letter reversals. Journal of Reading Behavior, 7(4):329–340, 1975. doi:10.1080/10862967509547152.

40. Marc H Bornstein, Charles G Gross, and Joan Z Wolf. Perceptual similarity of mirror images in infancy. Cognition, 6(2):89–116, 1978. doi:10.1016/0010-0277(78)90017-3.

41. James M Cornell. Spontaneous mirror-writing in children. Canadian Journal of Psychology/Revue canadienne de psychologie, 39(1):174, 1985. doi:10.1037/h0080122.

42. Stanislas Dehaene, Kimihiro Nakamura, Antoinette Jobert, Chihiro Kuroki, Seiji Ogawa, and Laurent Cohen. Why do children make mirror errors in reading? neural correlates of mirror invariance in the visual word form area. Neuroimage, 49(2):1837–1848, 2010. doi:10.1016/j.neuroimage.2009.09.024.

43. JE Rollenhagen and CR Olson. Mirror-image confusion in single neurons of the macaque inferotemporal cortex. Science, 287(5457):1506–1508, 2000. doi:10.1126/science.287.5457.1506.

44. Gordon C Baylis and Jon Driver. Shape-coding in it cells generalizes over contrast and mirror reversal, but not figure-ground reversal. Nature neuroscience, 4(9):937–942, 2001. doi:10.1038/nn0901-937.

45. Vadim Axelrod and Galit Yovel. Hierarchical processing of face viewpoint in human visual cortex. Journal of Neuroscience, 32(7):2442–2452, 2012. doi:10.1523/JNEUROSCI.4770-11.2012.

46. Tim C Kietzmann, Jascha D Swisher, Peter König, and Frank Tong. Prevalence of selectivity for mirror-symmetric views of faces in the ventral and dorsal visual pathways. Journal of Neuroscience, 32(34):11763–11772, 2012. doi:10.1523/JNEUROSCI.0126-12.2012.

47. Fernando M Ramírez, Radoslaw M Cichy, Carsten Allefeld, and John-Dylan Haynes. The neural code for face orientation in the human fusiform face area. Journal of Neuroscience, 34(36):12155–12167, 2014. doi:10.1523/JNEUROSCI.3156-13.2014.

48. Tim C Kietzmann, Anna L Gert, Frank Tong, and Peter König. Representational dynamics of facial viewpoint encoding. Journal of cognitive neuroscience, 29(4):637–651, 2017. doi:10.1162/jocn_a_01070.

49. Daniel D Dilks, Joshua B Julian, Jonas Kubilius, Elizabeth S Spelke, and Nancy Kanwisher. Mirror-image sensitivity and invariance in object and scene processing pathways. Journal of Neuroscience, 31(31):11305–11312, 2011. doi:10.1523/JNEUROSCI.1935-11.2011.

50. DI Perrett, MW Oram, MH Harries, R Bevan, JK Hietanen, PJ Benson, and S Thomas. Viewer-centred and object-centred coding of heads in the macaque temporal cortex. Experimental brain research, 86(1):159–173, 1991. doi:10.1007/BF00231050.

51. Nikos K Logothetis, Jon Pauls, and Tomaso Poggio. Shape representation in the inferior temporal cortex of monkeys. Current biology, 5(5):552–563, 1995. doi:10.1016/S0960-9822(95)00108-4.

52. Michael C. Corballis and Ivan L. Beale. The psychology of left and right. The psychology of left and right. Lawrence Erlbaum, Oxford, England, 1976.

53. Charles G Gross, David B Bender, and Mortimer Mishkin. Contributions of the corpus callosum and the anterior commissure to visual activation of inferior temporal neurons. Brain research, 131(2):227–239, 1977. doi:10.1016/0006-8993(77)90517-0.

54. Kaiming He, Xiangyu Zhang, Shaoqing Ren, and Jian Sun. Deep residual learning for image recognition. In Proceedings of the IEEE conference on computer vision and pattern recognition, pages 770–778, 2016. URL https://openaccess.thecvf.com/content_cvpr_2016/html/He_Deep_Residual_Learning_CVPR_2016_paper.html.

55. Umut Güçlü and Marcel AJ van Gerven. Deep neural networks reveal a gradient in the complexity of neural representations across the ventral stream. Journal of Neuroscience, 35(27):10005–10014, 2015. doi:10.1523/JNEUROSCI.5023-14.2015.

56. Martin Schrimpf, Jonas Kubilius, Ha Hong, Najib J. Majaj, Rishi Rajalingham, Elias B. Issa, Kohitij Kar, Pouya Bashivan, Jonathan Prescott-Roy, Franziska Geiger, Kailyn Schmidt, Daniel L. K. Yamins, and James J. DiCarlo. Brain-score: Which artificial neural network for object recognition is most brain-like? bioRxiv preprint, 2018. doi:10.1101/407007.

57. Martin Schrimpf, Jonas Kubilius, Michael J. Lee, N. Apurva Ratan Murty, Robert Ajemian, and James J. DiCarlo. Integrative benchmarking to advance neurally mechanistic models of human intelligence. Neuron, 108 (3):413–423, November 2020. doi:10.1016/j.neuron.2020.07.040.

58. Katherine R. Storrs, Tim C. Kietzmann, Alexander Walther, Johannes Mehrer, and Nikolaus Kriegeskorte. Diverse Deep Neural Networks All Predict Human Inferior Temporal Cortex Well, After Training and Fitting. Journal of Cognitive Neuroscience, 33(10):2044–2064, September 2021. doi:10.1162/jocn_a_01755.

59. Pinglei Bao, Liang She, Mason McGill, and Doris Y Tsao. A map of object space in primate inferotemporal cortex. Nature, 583(7814):103–108, 2020. doi:10.1038/s41586-020-2350-5.

60. Thomas Gerig, Andreas Morel-Forster, Clemens Blumer, Bernhard Egger, Marcel Luthi, Sandro Schönborn, and Thomas Vetter. Morphable face models-an open framework. In 2018 13th IEEE International Conference on Automatic Face & Gesture Recognition (FG 2018), pages 75–82. IEEE, 2018. doi:10.1109/FG.2018.00021.

61. Verena Willenbockel, Javid Sadr, Daniel Fiset, Greg O Horne, Frédéric Gosselin, and James W Tanaka. Controlling low-level image properties: the SHINE toolbox. Behavior research methods, 42(3):671–684, 2010. doi:10.3758/BRM.42.3.671.

62. Olga Russakovsky, Jia Deng, Hao Su, Jonathan Krause, Sanjeev Satheesh, Sean Ma, Zhiheng Huang, Andrej Karpathy, Aditya Khosla, Michael Bernstein, Alexander C. Berg, and Li Fei-Fei. ImageNet large scale visual recognition challenge. International Journal of Computer Vision (IJCV), 115(3):211–252, 2015. doi:10.1007/s11263-015-0816-y.

63. The MathWorks, Inc. Deep Learning Toolbox. Natick, Massachusetts, United State, 2019. URL https://www.mathworks.com/help/deeplearning/.

64. Adam Paszke, Sam Gross, Francisco Massa, Adam Lerer, James Bradbury, Gregory Chanan, Trevor Killeen, Zeming Lin, Natalia Gimelshein, Luca Antiga, Alban Desmaison, Andreas Kopf, Edward Yang, Zachary DeVito, Martin Raison, Alykhan Tejani, Sasank Chilamkurthy, Benoit Steiner, Lu Fang, Junjie Bai, and Soumith Chintala. Pytorch: An imperative style, high-performance deep learning library. In Advances in Neural Information Processing Systems, volume 32. Curran Associates, Inc., 2019. URL https://proceedings.neurips.cc/paper/2019/file/bdbca288fee7f92f2bfa9f7012727740-Paper.pdf.

65. A. Vedaldi and K. Lenc. MatConvNet – convolutional neural networks for MATLAB. In Proceeding of the ACM Int. Conf. on Multimedia, 2015. doi:10.1145/2733373.2807412.

66. Yann A LeCun, Léon Bottou, Genevieve B Orr, and Klaus-Robert Müller. Efficient backprop. In Neural networks: Tricks of the trade, pages 9–48. 2012. doi:10.1007/978-3-642-35289-8_3.

67. Kaiming He, Xiangyu Zhang, Shaoqing Ren, and Jian Sun. Delving deep into rectifiers: Surpassing human-level performance on imagenet classification. In Proceedings of the IEEE International Conference on Computer Vision (ICCV), December 2015. URL https://openaccess.thecvf.com/content_iccv_2015/html/He_Delving_Deep_into_ICCV_2015_paper.html.

